# YAP/TAZ and EZH2 synergize to impair tumor suppressor activity of TGFBR2 in non-small cell lung cancer

**DOI:** 10.1101/2020.07.10.197392

**Authors:** Federica Lo Sardo, Claudio Pulito, Andrea Sacconi, Etleva Korita, Marius Sudol, Sabrina Strano, Giovanni Blandino

**Author notes:** Corresponding authors: Sabrina Strano: e-mail address, Regina Elena National Cancer Institute, Via Elio Chianesi, 53-00144, Rome-Italy., Giovanni Blandino, e-mail address, Regina Elena National Cancer Institute, Via Elio Chianesi, 53-00144, Rome-Italy.

## Abstract

Lung cancer is the leading cause of cancer-related deaths, worldwide. Non–small cell lung cancer (NSCLC) is the most prevalent lung cancer subtype. YAP and TAZ have been implicated in lung cancer by acting as transcriptional co-activators of oncogenes or as transcriptional co-repressors of tumor suppressor genes. Previously we reported that YAP and TAZ regulate microRNAs expression in NSCLC. Among the set of regulated miRNAs, the oncogenic miR-25, 93, and 106b, clustering within the MCM7 gene were selected for further studies. We firstly identified Transforming Growth Factor-β (TGF-β) Receptor 2 (TGFBR2), a member of the TGF-β signaling, as a target of the miRNA cluster, which exhibited prognostic value because of its tumor suppressor activity. We found that YAP/TAZ-mediated repression of TGFBR2 occurs both: post-transcriptionally through the miR-106b-25 cluster and transcriptionally by engaging the EZH2 epigenetic repressor that we reported here as a novel target gene of YAP/TAZ. Furthermore, we document that YAP/TAZ and EZH2 cooperate in lung tumorigenesis by transcriptionally repressing a specific subset of tumor suppressor genes, including TGFBR2. Our findings point to YAP/TAZ and EZH2 as potential therapeutic targets for NSCLC treatment.

## 1. Introduction

Lung cancer continues to be the leading cause of cancer deaths, worldwide (1). Current treatments that are based on surgery, radiation, chemotherapy, laser, and photodynamic therapy result in the modest increase of the overall survival of the patients. The <u>N</u>on-<u>S</u>mall <u>C</u>ell <u>L</u>ung <u>C</u>ancer (NSCLC) still remains one of the most aggressive subtypes of lung cancer with the lowest survival rate (2). The development of targeted therapies and, more recently immunotherapy, have improved the overall outcome for lung cancer patients (3) (4). However, ultimately most of the patients either suffer adverse side effects of therapeutic interventions or develop resistance. Consequently then, these therapies are interrupted or terminated. Combinatorial treatments are intensely investigated now because they allow the use of lower doses of drugs, reducing side effects, and they could likely bypass the mechanisms of resistance. In the context of lung cancer, YAP and TAZ, the downstream effectors of the Hippo tumor suppressor pathway promote cell proliferation, survival, migration, invasiveness, and immune escape, which *in vivo* results in tumor development, progression and resistance to therapies (reviewed in (5)). This makes YAP and TAZ attractive therapeutic targets in lung cancer. By deciphering signaling mechanisms that are orchestrated by YAP and TAZ, we should get a better insight into the molecular pathology of lung cancer in general. Previously, we found that YAP/TAZ, and its preferred TEAD1 transcription factor, regulate the expression of the oncogenic cluster of miR-25, 93, and 106b, which is located within the transcript of the Mini-Chromosome Maintenance 7 (MCM7) gene (6). The regulation of miRNAs and long non-coding RNAs, in addition to that of coding genes, further expands the plethora of oncogenic mechanisms that can be orchestrated by YAP/TAZ. Recently we described the post-transcriptional inhibition of the p21 tumor suppressor gene as one of the mechanisms that underlie NSCLC. In the present work, we focused on TGFBR2, a novel potential target of the miR-106b-25 cluster. The TGFBR2 gene encodes for the <u>T</u>ransforming <u>G</u>rowth <u>F</u>actor-<u>β</u> (TGF-β) receptor 2, a member of the TGF-β signaling, which is involved in embryonic development, tissue homeostasis, and tumorigenesis. The TGFBR2 signals through the regulation of cell growth, differentiation, apoptosis, invasion, angiogenesis, and immune response (7). Upon ligand binding, the TGFBR2 forms a hetero-tetrameric complex with TGFBR1, which phosphorylates and activates receptor-regulated SMADs (<u>S</u>mall, <u>M</u>others <u>A</u>gainst <u>D</u>ecapentaplegic), namely SMAD2 and SMAD3. In turn, activated SMAD2 and SMAD3 associate with SMAD4, translocate into the nucleus, and engage transcription factors and co-activators to regulate gene expression program that is cell-context dependent (8). Generally, in cancer, the TGF-β pathway can be either: pro-oncogenic, inducing epithelial-mesenchymal transition (EMT), migration, invasion, and metastasis, or tumor-suppressive, by inducing growth arrest, apoptosis, and prevention of immortalization. The signaling outcome, however, is always tumor- and cancer stage-dependent (9).

Here, we show that TGFBR2 acts as a *bona fide* tumor suppressor and therefore it could be of prognostic value for NSCLC, especially at the early stages of the disease. YAP/TAZ signal to maintain low expression levels of TGFBR2 in NSCLC through at least two molecular mechanisms: post-transcriptionally (mediated by miR-106b-25 cluster) and transcriptionally (mediated by the epigenetic repressor <u>E</u>nhancer of <u>Z</u>este <u>H</u>omologue <u>2</u>, EZH2). We also find that YAP/TAZ/EZH2 cooperate in the repression of the TGFBR2 gene, as well as other tumor suppressor genes. Therefore, the combinatorial targeting of YAP/TAZ and EZH2 may represent a novel therapeutic strategy for controlling NSCLC.

## 2. Materials and Methods

### Cell culture and transfection

Human H1299, H1975, A459 and U293 cells were purchased from the American Type Culture Collection (ATCC, Manassas, VA) and routinely tested by PCR for mycoplasma contamination by using the following primers: Myco_fw1: 5′-ACACCATGGGAGCTGGTAAT-3′, Myco_rev1: 5′-CTTCATCGACTTTCAG ACCCAAGGCA-3′. A459 cells were grown in RPMI medium (Invitrogen, Carlsbad, CA) supplemented with 10% fetal bovine serum and Pen/Strep antibiotic at 37°C in a balanced air humidified incubator with 5% CO2. Lipofectamine RNAimax (Invitrogen) was used in accordance with the manufacturer’s instruction for transfection with siRNAs, LNA and miRNA mimics. siRNAs were used at the final amount of 300 pmol in 60 mm dish. List of siRNA used for functional in vitro experiments is given in Supplementary Materials and Methods. LNA inhibitors for miR-25, 93 and 106b (Exiqon, Vedbk, Denmark) were used at a final amount of 150 pmol in 60 mm dish. For mature miR-25, 93 and 106b overexpression, we used mirVana™ miRNA Mimic Negative Control #1 (Ambion) or hsa-miR-25-3p, hsa-miR-93-5p, or hsa-miR-106b-3p mirVana™ miRNA Mimic (Ambion) at final concentration of 5 nM. Plasmids were transfected with Lipofectamine 2000 (Invitrogen) in accordance with the manufacturer’s instruction at a final concentration of 1 µg in a 60 mm dish. Cells were collected 48-72h post transfection for subsequent analyses.

### Plasmids

The plasmid for EZH2 overexpression was obtained by cloning the EZH2 cDNA into pCDNA3 backbone with a myc-tag and was a kind gift from the laboratory of Dr. Daniela Palacios (Santa Lucia Foundation, Rome Italy). The plasmid for TGFBR2 3’UTR luciferase assay (Psi-check2-TGFBR2) was a kind gift from the laboratory of Dr. Stefen Wiemann (German Cancer Research Center (DKFZ), Heidelberg, Germany).

### Stable transfection

H1299 and H1975 cells were transfected using Lipofectamine 2000 with a pCDNA3-myc-EZH2 plasmid. 24 h after transfection, cells were diluted at 20–30% confluency and fresh medium with 1ng/µg G418 was added for a selection of stably transfected cells every 3–4 days. Cells were selected for 15-20 days and then they were grown in fresh medium containing 1 ng/µl G418, tested for correct EZH2 overexpression and expanded. For all experiments, cells were maintained in fresh medium containing 0,5 µg/µl G418.

### Clonogenic assay

Cells were transfected as indicated above and 48h-72h later they were detached and seeded at 500–1000 cells/well into 6-well or 12-well dishes. Fresh medium was added every 4 days. After 7–14 days, colonies were stained with crystal violet and counted.

### Pharmacological treatment and Chemical reagents

Dasatinib and Tazemetostat (EPZ-6438) were obtained from Selleck Chemicals (Houston, TX); To determine the IC50 of these drugs, lung adenocarcinoma cells were seeded in triplicate at a density of 1,000 cells/well. The following day, cells were treated with the drugs at increasing concentrations, and ATP lite assay (Promega) was performed after 72 hours of treatment. The dual drug studies (Dasatinib plus Tazemetostat) were performed in a similar manner with the doses indicated in the figures.

For colony assay upon treatment with Dasatinib and Tazemetostat, cells were grown for 9 days with or without Tazemetostat (added every 3-4 days) at different indicated doses, then they were seeded in triplicate at a density of 1,000 cells/well in a six-well multiwell. Every 3-4 days, fresh medium was added with or without dasatinib and Tazemetostat, alone or in combination, at the indicated doses. Colonies were stained with crystal violet and counted after 10–14 days.

### FACS cell cycle analysis

For cell cycle analysis, cells were collected 48-72h after interference, fixed in 70% ethanol and stored at −20°C (up to weeks). Fixed cells were treated with RNase at 1 mg/ml final concentration for 30 min at 37°C or overnight at 4°C before adding 5 mg/ml PI and analyzed with Guava Easycyte 8HT flow cytometer equipped with Guava Soft 2.1 (Millipore).

### Protein extracts and Western blot analysis

For the preparation of whole-cell lysates, cells were lysed in buffer with 50 mM Tris–HCl pH 7.6, 0.15 M NaCl, 5 mM EDTA, 1% Triton X-100 and fresh protease inhibitors. Extracts were sonicated for 10 + 15s at 80% amplitude and centrifuged at 12 000∼ rpm for 10 min to remove cell debris. For preparation of nuclear and cytoplasmic extracts, cells were lysed in a Cytoplasmic Extract (CE) buffer (10mM Tris-Cl pH 7.5, 60mM KCl, 1mM EDTA, 0,075% NP40, proteinase inhibitors) for 3 minutes on ice. The lysate was then centrifuged at 1500 rpm at 4°C for 4 minutes. The supernatant (Cytoplasmic Extract) was collected into a fresh tube. The pellet was washed three times in cold CE buffer without NP40 and lysed in Nuclear Extract (NE) buffer (20mM Tris-Cl pH 8.0, 420mM NaCl, 1,5 mM MgCl_2_, 0,2 mM EDTA, proteinase inhibitors) for 10 munites and sonicated. CE and NE extracts were then centrifuged at max speed for 10 minutes to pellet any residual nuclei. Protein concentrations were determined by colorimetric assay (Bio-Rad). Antibodies used for Western Blotting are listed in the Supplementary Materials and Methods.

### MiRNA and transcript analysis

Total RNA was extracted using TRIzol (Ambion) according to the manufacturer’s recommendations. For miR analysis, 30 ng RNA was retrotranscribed using the TaqMan microRNA Reverse Transcription Kit (Applied Biosystem) and Real-time PCR of miR expression was carried out in a final volume of 10 µl using TaqMan MicroRNA Assays (Applied Biosystems) and normalized on RNU48 and RNU49 as endogenous controls. We have chosen RNU48 and RNU49 because they were not modulated in our experimental conditions. TaqMan probes for miRNAs and RNU were purchased from Applied Biosystems. For gene transcript analysis, 1γ RNA was retrotranscribed using M-MLV reverse transcriptase (Invitrogen) following the manufacturer instructions. Real time PCR was performed into a final volume of 10 µl using Sybr Green PCR master mix, and normalized on GAPDH. All the real-time PCR assays were performed by using an Applied Biosystems® 7500 fast or *StepOne* Real-Time PCR Instruments. Each analysis was performed at least on three independent biological replicates. List of primers used for Real time PCR is given in Supplementary Materials and Methods.

### Chromatin Immunoprecipitation (ChIp)

ChIP-qPCR was performed as described previously (6) and the method is given in Supplementary Materials and Methods.

### Luciferase assay

Luciferase assay was performed as described previously (6) and the protocol is given in Supplementary Materials and Methods.

### Analysis of Differentially expressed genes

Deregulation of genes in different set of patient samples was assessed by two tailed student’s t test, and a false discovery rate procedure was performed to take into account multiple comparisons. The significance level was set to 5%. Analyses were performed by Matlab (The MathWorks Inc.). Association between pairs of genes was evaluated by calculating Pearson’s R correlation coefficient.

### Curves of OS and DFS

Curves of overall survival or disease-free survival in TCGA patients were evaluated by Kaplan–Meier method. Curves of patients with high and low signals were considered to establish statistical significance by using the logrank test. Analyses were performed by Matlab (The MathWorks Inc.). Disease-free survival includes both progression free survival and recurrence free survival.

See Supplementary Material and Methods for the list of antibodies used for western blotting, sequence of siRNA used for interference experiments and sequence of primers used for transcript analyses and for ChIp.

## 3. Results

### TGFBR2 acts as a tumor suppressor in lung cancer and is a direct target of miR-106b-25 cluster

To dissect mechanistically how MCM7 and its ‘hosted’ miR-106b-25 cluster elicit their oncogenic activities in lung cancer, we analyzed lung cancer genome atlas (TCGA), searching for transcripts that are inversely correlated to the miR cluster and therefore predict potential therapeutic targets for lung cancer. Indeed, we found a number of transcripts that code for proteins that signal in cancer pathways (6). From among them, we elected to focus on TGFBR2 that was inversely correlated with MCM7 in a robust way and hosted miRNAs with R Spearman ranging from −0,23 for miR-25 to −0,52 for MCM7 (**Fig. 1A**, Supplementary Fig. S1A). In cancer, the TGF-β pathway can be either pro-oncogenic or tumor-suppressive, depending on both the kind of tumor and its stage (9). Using the ‘FireBrowse Gene Expression’ computer platform (http://firebrowse.org) we analyzed transcript expression data of different cancer types of patient samples deposited in the TCGA dataset. We found that TGFBR2 expression is lower in tumor tissues when compared to non-tumor controls in several cancers, including <u>Lu</u>ng <u>Ad</u>enocarcinoma (LUAD), <u>Lu</u>ng <u>S</u>quamous cell <u>C</u>arcinoma (LUSC), the two main types of NSCLC (**Fig. 1B-C**, Supplementary Fig. S1B). Interestingly, in contrast to most of the analyzed tumor types, pancreatic adenocarcinomas and glioblastoma tissues exhibited higher levels of TGFBR2 expression, compared to non-tumoral controls. This data is in line with the notion of dual activity of TGFBR2: being either oncogenic or tumor-suppressive, depending on the cancer type (Supplementary Fig. S1B).

**Fig. 1.**
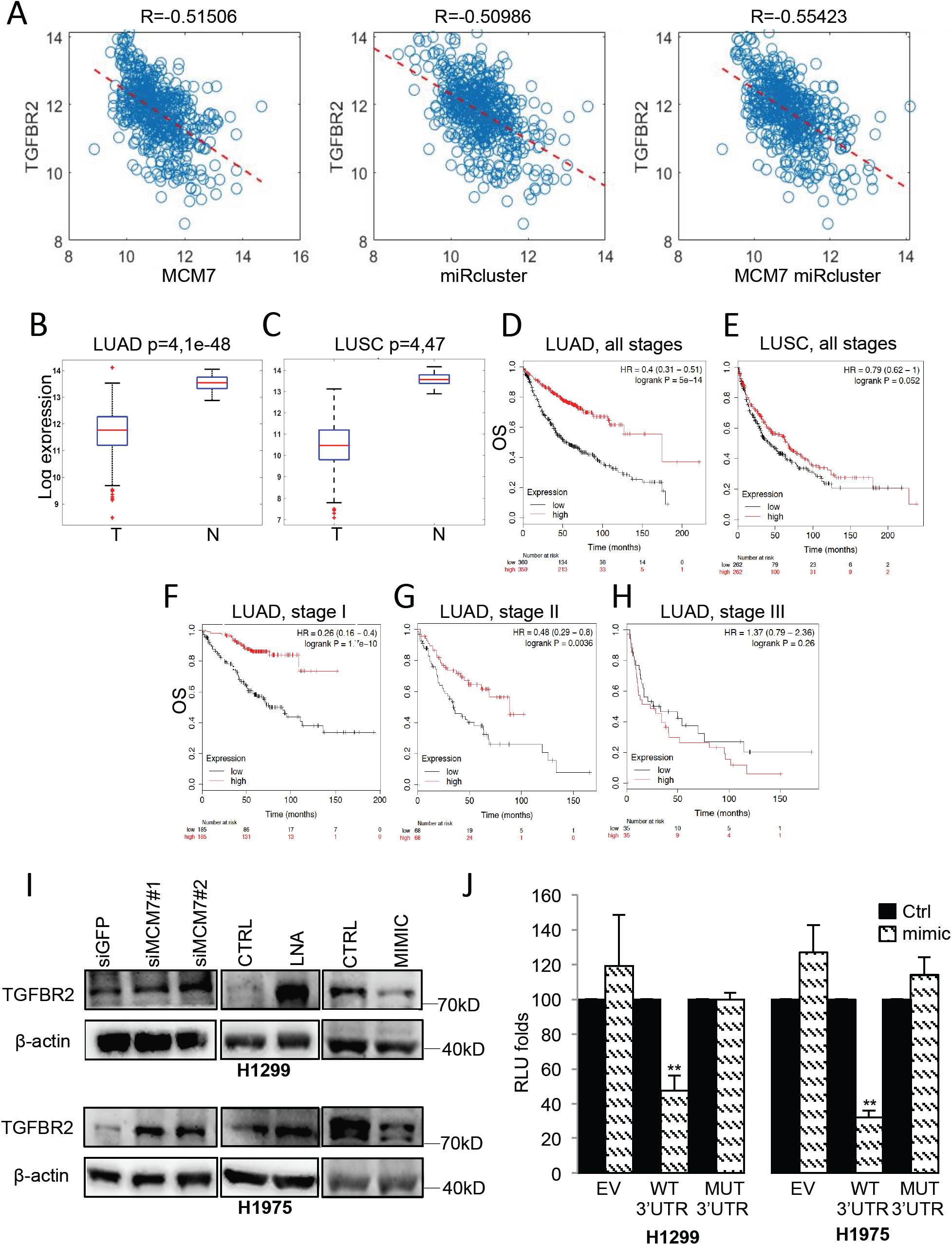
TGFBR2 is a direct target of the oncogenic miR-106b-25 cluster and exhibits prognostic value in NSCLC. **A**, Dot plots showing the correlation between TGFBR2 transcript and MCM7 (left panel), TGFBR2 and the miR106b-25 cluster (mid panel), and between TGFBR2 and MCM7/miR cluster together (right panel) in Lung Adenocarcinoma (LUAD) patients from the TCGA. **B**-**C**, Boxplot showing the expression of TGFBR2 transcript in tumoral (T) compared to non-tumoral (N) tissues in lung adenocarcinoma (LUAD) patients (**B**) and lung squamous cell carcinoma (LUSC) (**C**) from the TCGA dataset. Number of LUAD samples: 510T, 58N. Number of LUSC samples: 501T, 51N. **D-H**, Kaplan-Meier (KM) survival of lung adenocarcinoma (**D**) and squamous cell carcinoma patients (**E**), and KM of adenocarcinoma patients stratified for tumor stage (**F-H**), with high or low expression of TGFBR2. The number of patients is indicated below the plots. I, Western blot analysis for TGFBR2 expression upon depletion of MCM7 (left panels), upon either transfection with LNA inhibitors (mid panels) or mimics (right panels) for miR-25, 93, and 106b in H1975 cells (upper panels) and H1299 (lower panels). β-actin expression was used for equal protein loading. **J**, Luciferase assay of the psiCHECK2-TGFBR2-3’UTR reporter wild type or mutated in the seed sequence recognized by miR-93 and 106b in H1299 transiently co-transfected either with control mimic or mimic for miR-93 and miR-106b. Data are compared to signals obtained from cells co-transfected with the same mimic and psiCHECK2 Empty Vector as a control. Firefly luciferase was used to normalize the Renilla luciferase. All the experiments have been performed in triplicate. Data are presented as mean ± SEM. Two-tailed t-test analysis was applied to calculate the P values. *p<0,05; **p<0,01;***p<0,001

Based on the median level of TGFBR2 transcript (10) the stratification of lung cancer patients from TCGA was performed. The patients were categorized into either a high- or low-expressing group. The data revealed that patients with a lower expression of TGFBR2 exhibited a shorter OS (overall survival) than those with a higher level of TGFBR2. This was even more evident in LUAD than in LUSC patients (**Fig. 1D-E**). Stratification of LUAD patients based on the stage of the tumor indicated that the association of low TGFBR2 expression with shorter OS was stronger in early-stage (stage I) patients than in those in advanced stages (stages II and III) (**Fig. 1F-H**, Supplementary Fig. S1C) (9).

At the cellular level, we found that either the depletion of MCM7 transcript or the transfection of the locked-nucleic-acid-based-construct (LNA) that suppressed the endogenous miR-25/93/106b cluster led to the increase in the levels of the TGFBR2 transcript and its protein in two representatives NSCLC cell lines: H1299 and H1975 (Fig. **1I**, Supplementary Fig. S1D). Conversely, ectopic expression of synthetic miRNA mimics reduced the expression of TGFBR2, when compared to control cells (**Fig. 1J**).

Importantly, we demonstrated the direct binding of miR-93 and miR-106b to the TGFBR2-3’UTR through luciferase reporter assays. H1299 and H1975 cells were co-transfected with miRNAs mimics and the psiCHECK2 plasmid that contained the TGFBR2-3’UTR downstream of the luciferase gene. The cells exhibited reduced luciferase activity when compared to cells co-transfected with the same vector and control mimic (**Fig. 1J**). This effect was not observed in cells transfected with an empty vector or when the TGFBR2-3’UTR was mutated in the cognate sequence recognized by miR-93 and miR-106b (**Fig. 1J**, Supplementary Fig. S1E). Collectively, these findings document the tumor-suppressive activity of TGFBR2 and its post-transcriptional regulation by the oncogenic miR-25/93/106b cluster in cell culture models of lung cancer.

### YAP and TAZ depletion affects the tumor suppressor TGF-β signaling

We previously showed that YAP and TAZ transcriptionally regulate MCM7 and the miRs that are located within the MCM7 gene (6). Herein, MCM7 and its hosted miRs are shown as negative regulators of TGFBR2 (**Fig. 1** and Supplementary Fig. S1A, D). Thus we sought to investigate whether the depletion of YAP/TAZ was able to affect the level of TGFBR2. We found that YAP/TAZ depletion led to the up-regulation of both TGFBR2 transcript and protein as a consequence of reduced expression of MCM7 and its miR cluster, as well (**Fig. 2A-B**, Supplementary Fig. S1F-G). Reassuringly, the ectopic expression of miR25, 93, and 106b upon YAP/TAZ depletion partially rescued this effect as well as the ability of H1299 and H1975 cells to form colonies (**Fig. 2C-D**, Supplementary Fig. S1H-K). The increased expression of TGFBR2 in siYAP/TAZ cells correlated with an increase of total SMAD2 and SMAD3 in the cytoplasm and of pSMAD3 in the nucleus (**Fig. 2E-F**, Supplementary Fig. S2A-B). TGF-β-induced tumor suppressor response caused a cell accumulation at the G1/S transition that paired with increased expression of the p21 and p15 tumor suppressors and reduced expression of the CDK activator cdc25A, respectively. This occurred through p53-independent mechanisms involving SMADs and Sp1 (11) (12) (13). In agreement with these items of evidence, a depletion of SMAD2/3/4 partially rescued the colony-forming potential of siYAP/TAZ cells (**Fig. 2G**, Supplementary Fig. S2C-G) with a partial rescue of p21 repression and cdc25A expression (**Fig. 2F**, Supplementary Fig. S2B). These results suggest that the signaling by TGFBR2 tumor suppressor that is orchestrated by the YAP/TAZ/MCM7 axis does involve SMADs.

**Fig. 2.**
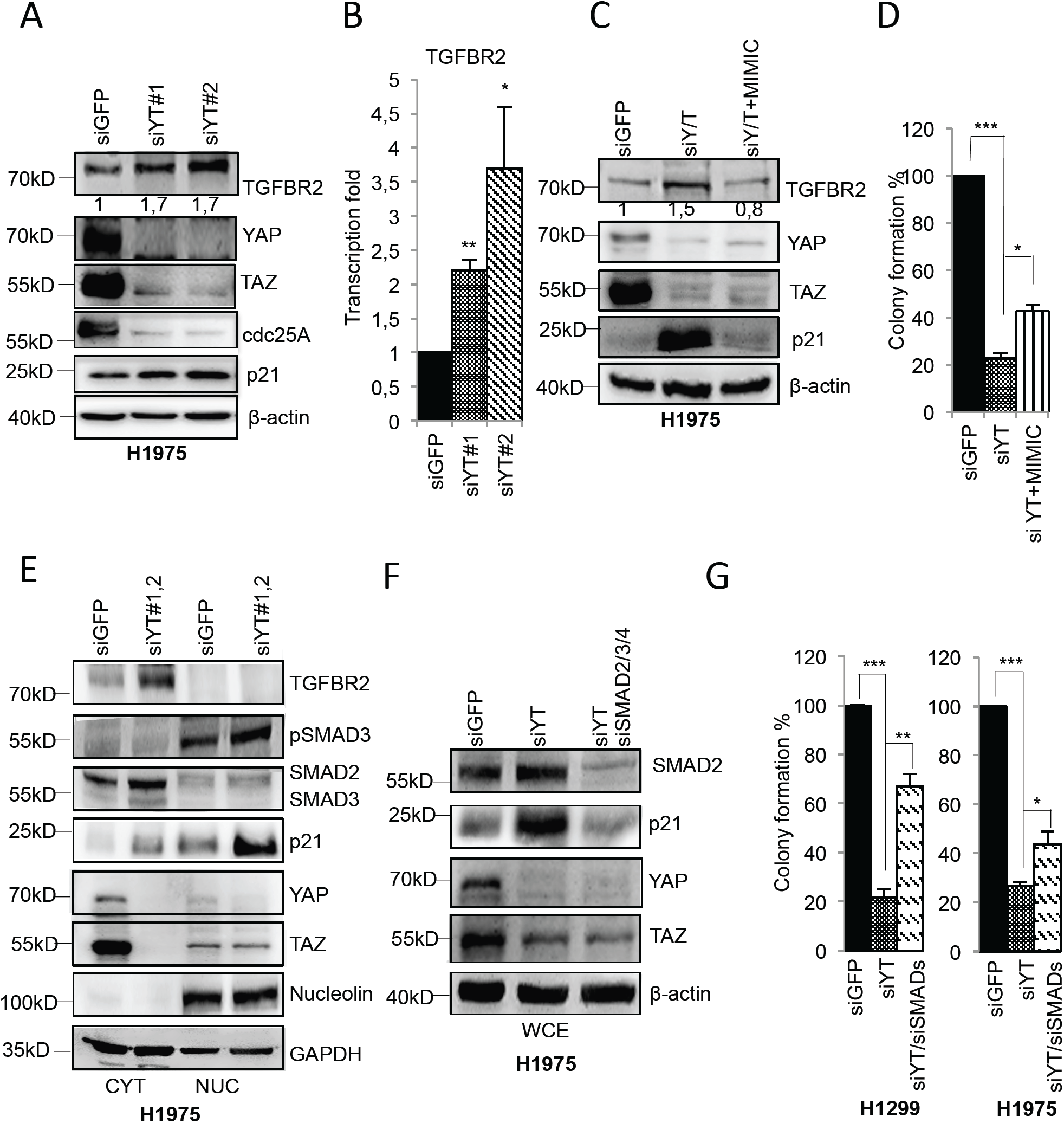
YAP/TAZ control TGFBR2 expression acting upstream to the miR cluster. **A-B**, Western blot analysis for expression of the indicated proteins normalized to β-actin (**A**) and quantification by real time-PCR of TGFBR2 transcript normalized to GAPDH (**B**) in H1975 cells depleted for YAP and TAZ proteins compared to control cells. The experiments have been performed in triplicate. **C**, Western blot analysis of the indicated proteins, normalized to β-actin in H1975 upon the interference of YAP and TAZ with or without mimic miR-93 and 106b. **D**, Quantification of colony formation of H1975 cells upon YAP/TAZ interference with or without concomitant transfection with mimic miR-93 and 106b, compared to controls. Data are presented as mean ± SEM. Two-tailed t-test analysis was applied to calculate the P values. *p<0,05; **p<0,01;***p<0,001. **E**, Western blot analysis of nucleo-cytoplasmic extracts from H1975 cells showing the abundance of the indicated proteins upon YAP/TAZ interference compared to siGFP control cells. Nucleolin and β-actin were used as a nuclear and cytoplasmic loading control, respectively. **F**, Western blot analysis of whole-cell extracts (WCE) of the indicated proteins normalized to β-actin in H1975 cells upon the interference of YAP and TAZ with two different combinations of alternative siRNA. **G**, Quantification of colony formation in H1299 and H1975 cells upon depletion of YAP/TAZ with or without interfering with SMAD2/3/4 expression compared to control cells.

### EZH2 mediates YAP transcriptional repression of TGFBR2

Our findings document that the increase of the TGFBR2 transcript is higher upon YAP/TAZ interference (**Fig. 2B**, Supplementary Fig. S1G) than upon miRNA depletion (Supplementary Fig. S1D). This suggests that YAP/TAZ may repress TGFBR2 through additional mechanisms. We noticed that the <u>E</u>nhancer of <u>Z</u>este <u>H</u>omologue <u>2</u> (EZH2), the enzymatic component of the <u>P</u>olycomb <u>R</u>epressive <u>C</u>omplex <u>2</u> (PRC2), was previously found as an epigenetic repressor of TGFBR2 in lung cancer (14). EZH2 is overexpressed in diverse cancer types, among them is lung cancer (Fig. 3a, Additional file 3: Figure S3a) in which EZH2 contributes to the increase in tumor growth, metastatic potential and therefore results in poor outcome and resistance to therapies. This occurs through the aberrant repression of tumor-suppressor genes, which is mediated by the enzymatic three-methylation of Lysine 27 of histone H3 (H3K27me3) (15). Here we found that EZH2 depletion reduced the colony-forming potential and affected the expression of cell cycle-related genes in both H1299 and H1975 lung cancer cell lines (**Fig. 3B-C**. Supplementary Fig. S3B). An inverse correlation between EZH2 and TGFBR2 transcripts was found in the TCGA lung cancer dataset (**Fig. 3D**). Interestingly, we observed that EZH2 depletion released the expression of TGFBR2 transcript in NSCLC cells (**Fig. 3E**, Supplementary Fig. S3C). It has been previously reported that YAP and TAZ depletion affected PRC2/EZH2 signature genes in melanoma cells (16). Furthermore, YAP favored the recruitment of the PRC2 complex onto the promoter of the GDF15 gene in breast cancer cells, whose transcriptional repression promoted metastasis (17). Moreover, YAP was shown to regulate the E2F1 transcription factor (18,19), and to cooperate with E2F1 in the regulation of many cell cycle-related genes (20). E2F1, in turn, was shown to regulate EZH2 expression through the binding onto its promoter (21). Therefore, we asked whether YAP and TAZ could regulate either EZH2 expression or cooperate with EZH2 in NSCLC. Interestingly, both EZH2 and YAP/TAZ depletion promoted the expression of two direct EZH2 target genes (22,23) such as PUMA and p16 in lung cancer cell lines (**Fig. 3C**, Supplementary Fig. S3D). Moreover, ectopic expression of EZH2 in lung cancer cells concomitantly depleted of YAP and TAZ proteins partially rescued their colony-forming ability and also partially reversed the expression of TGFBR2 (**Fig. 3F-I**). Notably, we found that YAP/TAZ interference in H1299 and H1975 cells reduced both EZH2 transcript and protein expression (**Fig. 4A**, Supplementary Fig. S3E, right panel). The other PRC2 complex components, EED and SUZ12, were not affected at the transcriptional level but were reduced at the protein level (Supplementary Fig. S3F-G). This might be due to the reduced stability of the PRC2 complex upon reduced expression of EZH2. This effect was more evident for SUZ12 (Supplementary Fig. S3F). In line with these observations, lower levels of SUZ12 protein were also observed upon EZH2 interference (Supplementary Fig. S3H-I). In sum, these findings indicate that YAP and TAZ are regulating the abundance of EZH2 in lung cancer cell lines.

**Fig. 3.**
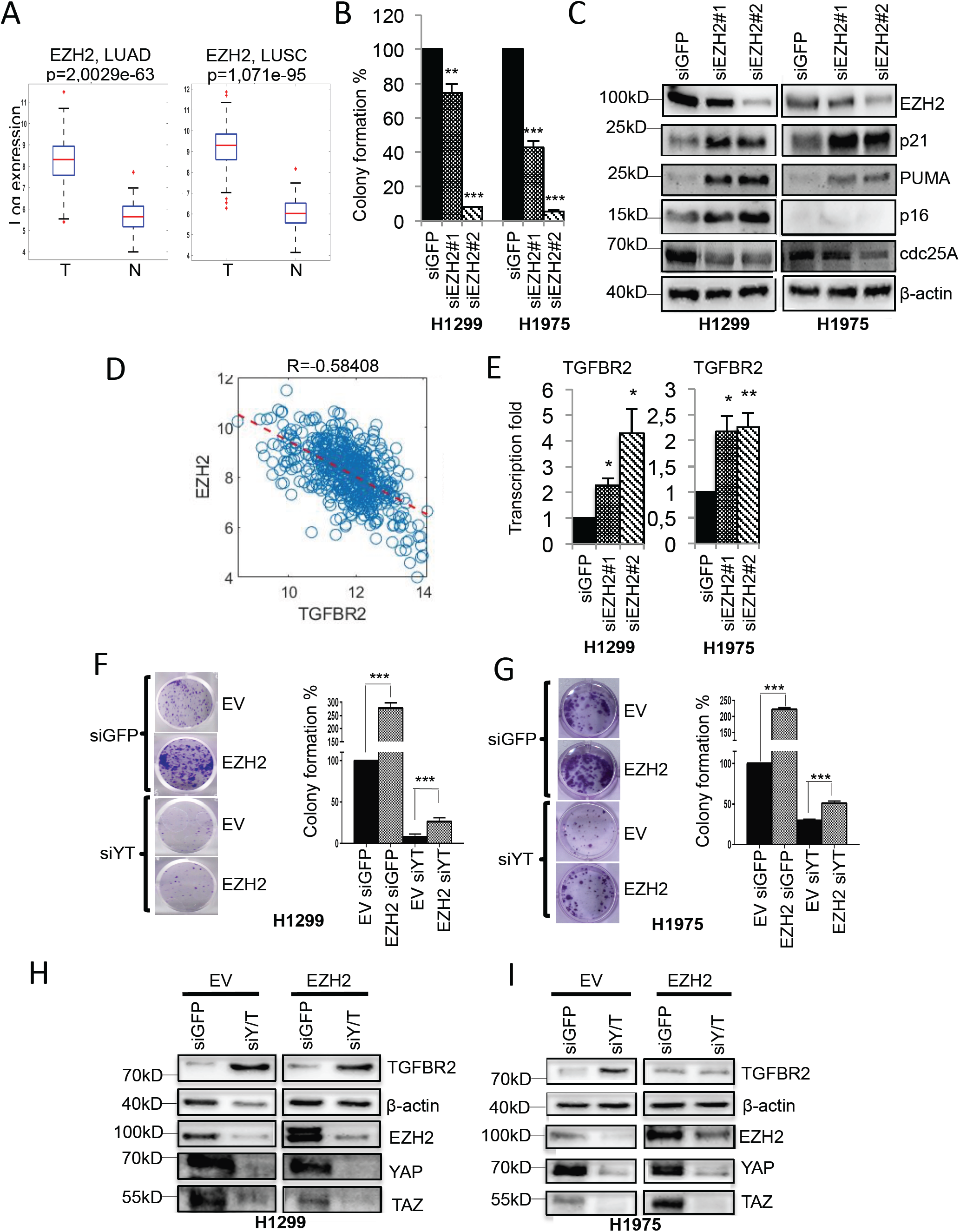
YAP and TAZ inhibit transcriptionally TGFBR2 expression through the oncogenic EZH2 repressor. **A**, Boxplot of the abundance of EZH2 in tumoral (T) compared to non-tumoral (N) tissues in lung adenocarcinoma patients (LUAD, left) and lung squamous cell carcinoma (LUSC, right) from the TCGA. The number of LUAD samples: 510T, 58N. The number of LUSC samples: 501T, 51N. **B**, Quantification of colony formation in H1299 (left) and H1975 (right) upon EZH2 depletion compared to control cells. **C**, Western blot analysis of the indicated proteins normalized to β-actin in H1299 (left) and H1975 cells (right) upon depletion of EZH2 protein. **D**, Dot plot showing the correlation between TGFBR2 and EZH2 transcripts in lung adenocarcinoma patients. **E**, RT-PCR quantification of TGFBR2 transcript upon depletion of EZH2 in H1299 (left) and H1975 (right). **F-G**, Representative images of cell colony formation assay (left panels) and quantification of relative colony formation (right panels) in H1299 (**F**) and H1975 (**G**) treated with siRNA against YAP/TAZ with or without concomitant overexpression of EZH2 compared to cells transfected with siGFP and empty vector as a control. Data are presented as mean ± SEM of three technical replicates of one representative experiment. Two-tailed t-test analysis was applied to calculate the p values. The experiment has been performed in triplicate. **H-I**, Western blot analysis of the indicated proteins normalized to β-actin in H1299 (**H**) and H1975 (**I**).

**Fig. 4.**
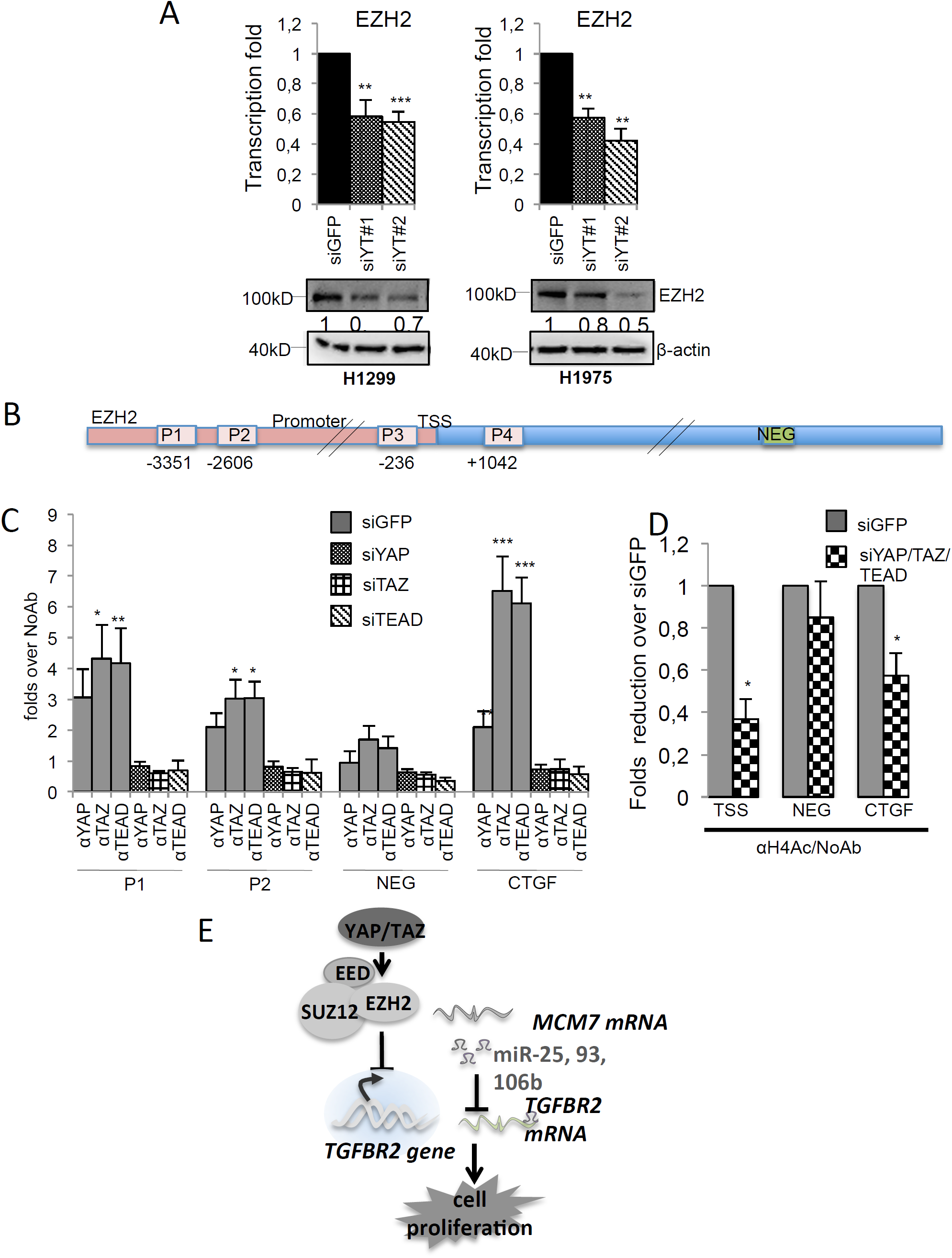
YAP/TAZ regulates EZH2 expression. **A**, RT-PCR quantification (upper panel) and western blot analysis (lower panel) of EZH2 transcript and protein, respectively, in H1299 (left) and H1975 cells (right) upon YAP and TAZ interference with two combinations of alternative siRNAs. **B**, Schematic representation of the EZH2 locus and the rgions containing the putative consensus for TEAD (P1, P2, P3, P4) with their relative position respect to TSS, and with the region used as a negative control of YAP/TAZ/TEAD binding (NEG). **C**, Fold enrichment of YAP, TAZ, and TEAD1 proteins onto the indicated sites of EZH2 locus in H1299 cells depleted for YAP, TAZ, and TEAD1 compared to control cells. CTGF promoter was used as a positive control while an intronic region of the EZH2 locus was used as a negative control. Data are presented as mean ± SEM of at least three biological replicates. For each antibody, fold enrichment was calculated over no antibody control. **D**, Chromatin Immunoprecipitation analysis of the abundance of H4Ac onto the indicated genomic sites (TSS, EZH2 NEG, and CTGF) in cells simultaneously depleted for YAP/TAZ/TEAD1 compared to siGFP as control. Fold enrichment was calculated over no antibody control and then normalized to the siGFP control that was adjusted to 1. The experiments were performed in triplicate. Two-tailed t-test analysis was applied to calculate the P values. *p<0,05; **p<0,01;***p<0,001. **E**, Working model indicating that YAP and TAZ downregulate TGFBR2 expression through miR-25,93 and 106b (post-transcriptionally) and the transcriptional repressor EZH2 (transcriptionally) in NSCLC.

### YAP/TAZ/TEAD is recruited onto EZH2 promoter

Using the “LASAGNA Search” software we analyzed genomic sequences for the TEAD-binding sites in the EZH2 promoter. We found four potential binding sites in proximity of the transcription start site (TSS) of the EZH2 promoter (P1 and P2, P3, P4 **Fig. 4B**). The analysis of “UCSC Genome Browser” indicated that P3 and P4 sites could be occupied by E2F1 and TEAD4 transcription factors (Supplementary Fig. S4A, red boxes). Chromatin Immunoprecipitation (CHIP) assays revealed that YAP, TAZ, and TEAD were recruited onto the EZH2 promoter regions in H1299 cells (**Fig. 4C**, Supplementary Fig. S4B). As expected, no specific enrichment was detected upon YAP, TAZ, or TEAD depletion. Also, in a control assay, we did not observe any complexes on an arbitrary intronic region that did not contain TEAD-binding sites (**Fig. 4B-C**, Supplementary Figs. S3E, S4A, green box). Moreover, the acetylation of histone H4, a mark of active transcription, was strongly reduced at the TSS upon YAP/TAZ/TEAD interference (**Fig. 4D**, Supplementary Fig. S3E). These items of evidence confirm that YAP, TAZ, and TEAD regulate EZH2 in NSCLC at the level of transcription. Collectively, our findings suggest that YAP/TAZ and the PRC2 complex cooperate to inhibit the expression of TGFBR2 through transcriptional mechanisms (**Fig. 4E**).

### YAP/TAZ and EZH2 co-repress tumor suppressor genes in NSCLC

We performed single and combined depletion of YAP/TAZ and EZH2 in H1299 and H1975 lung cancer cell lines. Combined interference was more effective than single interference in the inhibition of cell cycle progression, as measured through the expression of the cell cycle regulator cdc25A (**Fig. 5A**, Supplementary Fig. S4C) and through cell cycle profile (**Fig. 5B**, Supplementary Fig. S4D). In addition, YAP/TAZ and EZH2 depletion synergistically affected colony formation of lung cancer cell lines (**Fig. 5C**, Supplementary Fig. S4E).

**Fig. 5.**
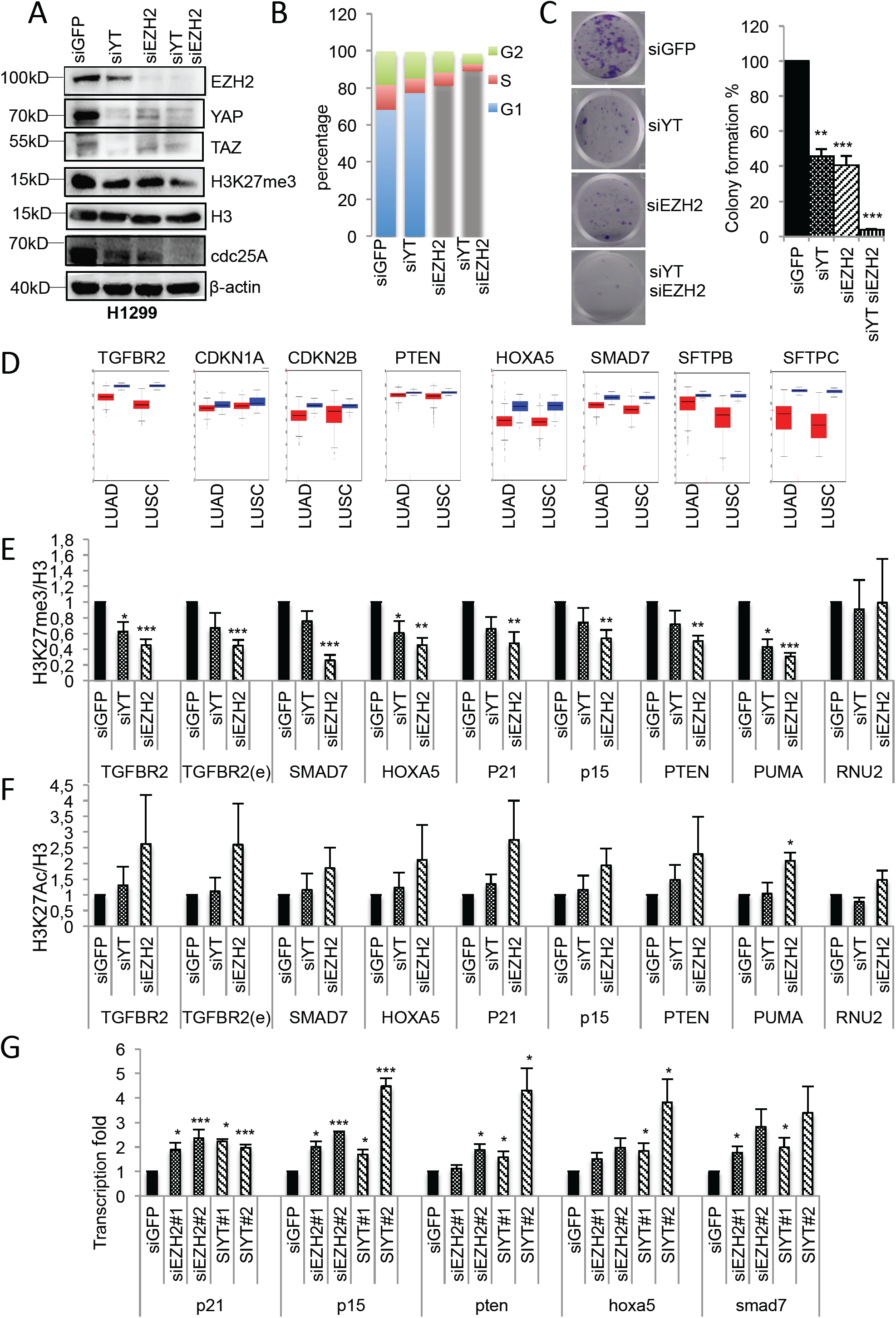
YAP/TAZ and EZH2 synergistically affect the cell growth of lung cancer cell lines. **A**, Western blot analysis of the indicated proteins in H1299 cells upon depletion of YAP/TAZ and EZH2, either alone or in combination, compared to control cells. **B-C**, Cells (%) in G1, S, and G2 phases (**b**) and colony quantification (**c**) of H1299 cells, upon depletion of YAP/TAZ or EZH2, alone or in combination, respect to control cells. **D**, Boxplots representing the expression profile of the indicated transcripts in LUng Adeno Carcinoma (LUAD) and Lung Squamous cell Carcinoma (LUSC) from patient samples of TCGA dataset as obtained from the FireBrowse Gene Expression Viewer. **E-F**, Chromatin Immunoprecipitation fold enrichment of H3K27me/H3 (**E**) and H3K27Ac/H3 (**F**) on the indicated loci. TGFBR2(e) indicates the enhancer region shown in figure S5a. For each antibody, fold enrichment was calculated over no antibody control and normalized to the siGFP control that was adjusted to 1. Experiments were performed in triplicate. Two-tailed t-test analysis was applied to calculate the P values. *p<0,05; **p<0,01;***p<0,001. (G) Real-time PCR analysis of the indicated transcripts in H1299 cells upon depletion of either YAP/TAZ or EZH2 proteins.

To understand whether co-repression of tumor-suppressor genes as for TGFBR2 could be a broad oncogenic activity mediated by YAP/TAZ/TEAD/EZH2, we searched the published literature for other genes that could be potentially repressed by both YAP/TAZ/TEAD and EZH2 in NSCLC or in other cell lines or models (**Table 1**). All of these genes were found to be associated with tumor suppression or cell differentiation and their aberrant repression was associated with tumorigenesis or stemness (6,24-30).

Using the “Firebrowse” (http://firebrowse.org) search platform, we found that genes listed in **Table 1** were down-regulated in tumors compared to non-tumoral tissues, both in lung adenocarcinomas and squamous cell carcinomas (**Fig. 5D**). Furthermore, by employing “Cistrome Data Browser” (http://cistrome.org), we revealed that YAP and TEAD4 factors as well as the H3K27me3 histone methylation signature occupied and were mapped, respectively, on the regulatory elements of these genes in lung cancer cell lines as well as in the IMR90 lung foetal cell line (Supplementary Fig. S5A-F). Interestingly, CHIP assays showed that the depletion of YAP/TAZ and EZH2 reduced the enrichment of H3K27me3 on these targets (**Fig. 5E**). These results were robust and statistically significant upon EZH2 depletion, while a general trend was seen upon YAP/TAZ interference (**Fig. 5E**). Interestingly, the enrichments in the H3K27Ac histone signature showed an opposite trend when compared to the H3K27me3 histone signature upon either YAP/TAZ or EZH2 depletion. These results revealed a common regulatory switch from repression (methylation) towards activation (acetylation) of the targeted loci (**Fig. 5F**). This effect was specific and was not observed for the U2 snRNA gene (RNU2), a constitutively transcribed locus that was used as a control (**Fig. 5E-F**). The total level of H3 was not affected in all of the analyzed targets (Supplementary Fig. S5G). Accordingly, the de-repression of the analyzed transcripts (**Table1**) was seen upon both YAP/TAZ and EZH2 depletion (**Fig. 5G**, Supplementary Fig. S5H).

**Table1.**
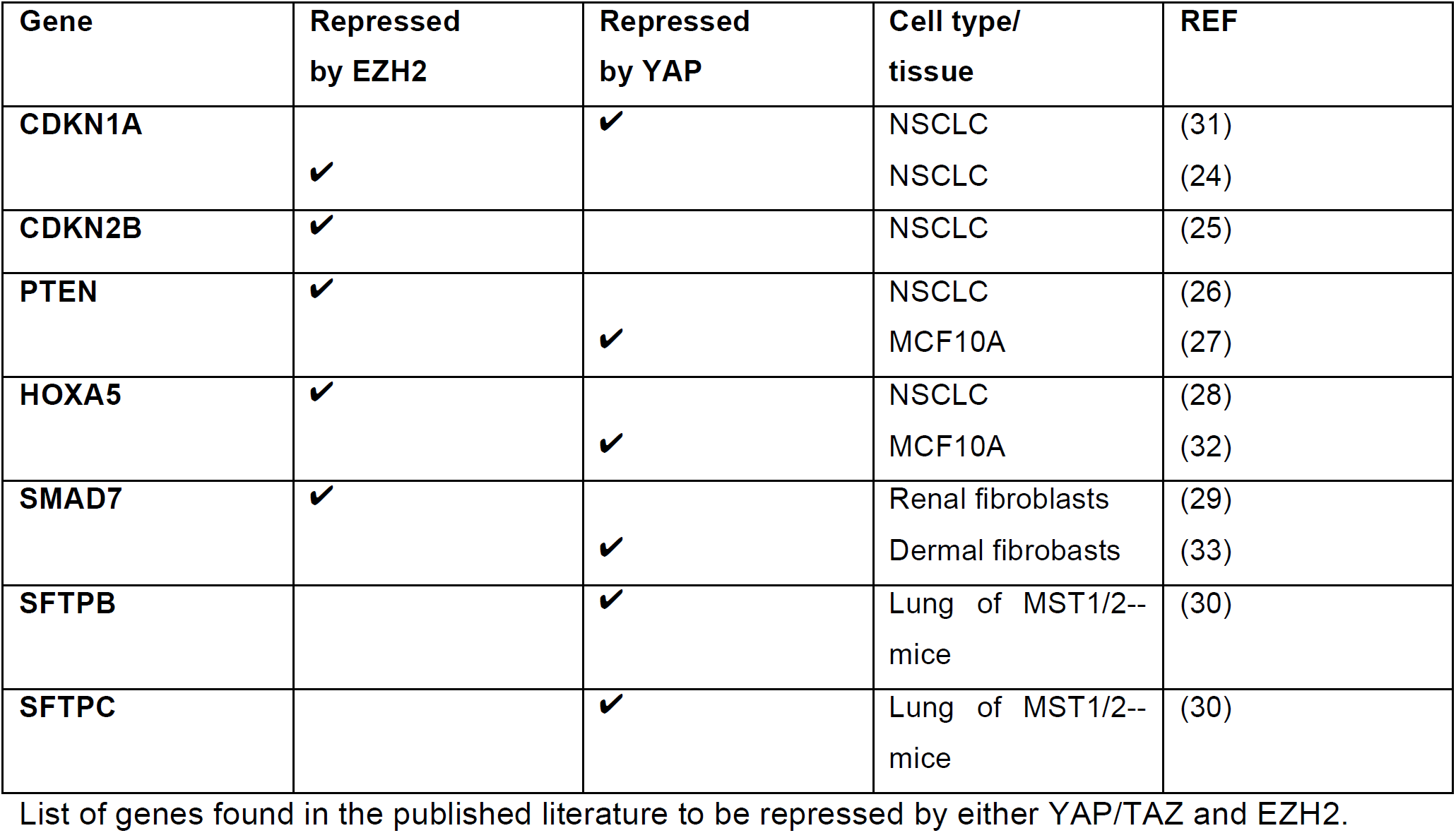
Genes repressed by YAP and EZH2 IN NSCLC.

Collectively, these findings indicate that co-repression of tumor suppressor genes by the concerted action of YAP/TAZ and EZH2 might play a broad role in lung tumorigenesis.

### Pharmacological targeting of YAP/TAZ and EZH2 affects synergistically lung cancer cell survival

We treated both H1299 and H1975 lung cancer cell lines with Dasatinib and Tazemostat (EPZ-6438) to pharmacologically inhibit YAP/TAZ (34) and EZH2 (35) respectively. Dasatinib treatment affected the transcriptional activity of YAP and TAZ in a dose-dependent manner. This was shown by a reduced expression of their well-known targets, such as CTGF and ANKRD1 (36) as well as the newly characterized target EZH2. However, TGFBR2 and p21 transcripts were de-repressed (Supplementary Fig. S6A, B). In addition, the expression of the cell cycle regulators cdc25A and c-myc was diminished, indicating a delayed cell cycle progression (Supplementary Fig. S6A, B). Tazemetostat treatment inhibited the enzymatic activity of EZH2 as shown by the reduction of H3K27me3 signatures globally, and it de-repressed PRC2 target genes in a dose and time-dependent manner (Supplementary Fig. S7A, B). This effect was more pronounced in H1299 than in H1975 cells (Supplementary Fig. S7A). In fact, H1975 were more resistant to Tazemetostat, as shown by the IC50 value measured through the “ATPlite” assay (Supplementary Fig. S8A, B). Some genes showed a more pronounced de-repression at longer time points, probably because of the slow kinetics of H3K27me3 turnover, as was shown previously (Supplementary Fig. S7A, B) (37). Functionally, the combination therapy with Dasatinib and Tazemetostat acted synergistically in affecting the colony formation and cell growth of H1299, H1975, as well as of A549 lung cancer cell line that was previously shown to be resistant to Dasatinib (38) (39) (**Fig. 6A-E**, Supplementary Fig. S8C-E). This effect was more evident in H1299, as observed at a very low dose of Tazemetostat (2µM) and with low doses of Dasatinib (0,025 µM) (**Fig. 6A-B**, Supplementary Fig. S8C). In the resistant A549 cells, increasing doses of Dasatinib up to 0,1µM in the absence of Tazemetostat did not reduce the number, but only the size of the colonies (**Fig. 6E**, lower panel, Supplementary Fig. S8E, left panel). However, the addition of Tazemetostat (2 µM) sensitized cells to Dasatinib and reduced both the number and size of the colonies (**Fig. 6E**, Supplementary Fig. S8E). While low amounts of Dasatinib alone, or Tazemetostat (2µM) alone, did not cause de-repression of TGFBR2, p21 or SMAD7 transcripts, the combination of the two drugs de-repressed these transcripts (to different extend, though) in both H1299 and H1975 cell lines (Supplementary Fig. S9A, B). In summary, these findings may provide a basis for testing the combination of Dasatinib and Tazemetostat as a therapeutic modality for the treatment of NSCLC patients (**Fig. 6F**).

**Fig. 6.**
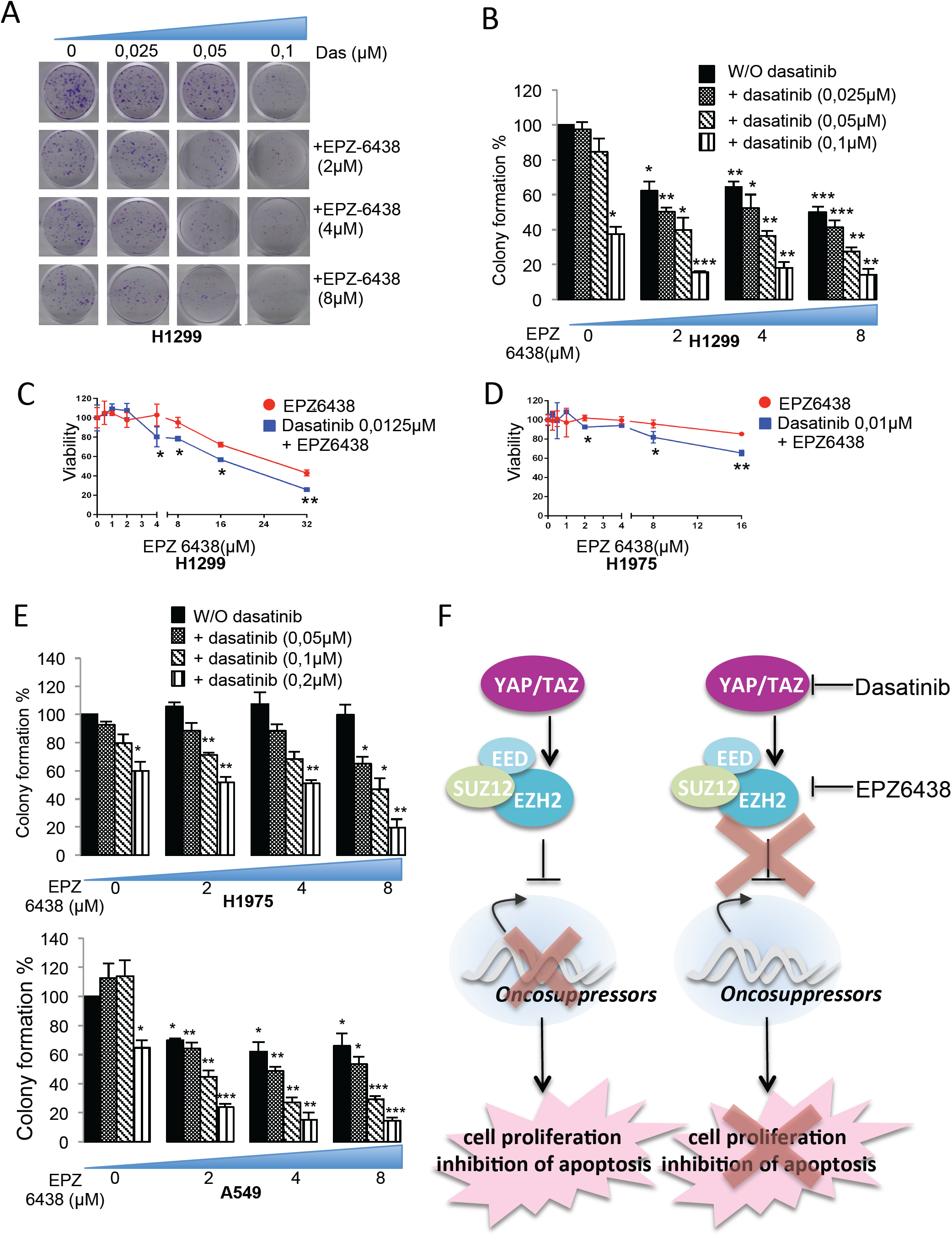
Low doses of Dasatinib with Tazemetostat synergistically affect cell proliferation. **A-B**, Representative images (**A**) and quantification (**B**) of colony formation upon treatment with different combinations of dasatinib and Tazemetostat (EPZ-6438) at the indicated doses in the H1299 cell line. **C-D**, Viability of H1299 cells (**C**) and H1975 cells (**D**) as measured with ATPlite assay after 72h treatment with a fixed dose of dasatinib and growing doses of Tazemetostat. **E**, Quantification of colony formation upon treatment with different combinations of Dasatinib and Tazemetostat in H1975 (upper panel) and A549 cells (lower panel). **F**, Schematic model of the oncogenic role of the YAP/TAZ/EZH2 axis which contribute to the aberrant proliferation of lung cancer cell lines. Pharmacological inhibition of YAP and EZH2 impairs the repression of tumor suppressor genes thereby reducing cell proliferation of lung cancer cell lines.

## 4. Discussion

We report here that YAP and TAZ elicit, at least in part, their oncogenic roles in NSCLC through the negative regulation of the tumor-suppressor “arm” of the TGF-β signaling pathway. It has been previously reported that YAP and TAZ crosstalk with the TGF-β pathway either by cooperating with its pro-tumorigenic signaling (i.e., induction of EMT) (40) or antagonizing its pro-apoptotic/tumor-suppressor activity. YAP prevents TGF-β1-mediated apoptosis without affecting EMT in normal mouse mammary epithelial cells (41). Similarly, YAP and TAZ promote TGF-β-induced EMT by inhibiting the TGF-β tumor suppression activity in breast cancer (42). Herein, we also show an antagonism between YAP/TAZ-oncogenic and TGF-β/SMAD-tumor-suppressive activities in NSCLC cells through the inhibition of TGFBR2. This inhibition is elicited through the aberrant activation of two oncogenic loci: EZH2 that represses TGFBR2 transcriptionally (14) and MCM7 that harbors the oncogenic cluster of miR106b-25. That mRNA cluster represses TGFBR2 at the post-transcriptional level. Clinically, lower expression of TGFBR2 in tumor tissues of lung cancer patients associates with poorer prognosis. This appears even more evident in the early stages of the disease.

Interestingly, the cooperation between YAP/TAZ and EZH2 to maintain the expression of tumor-suppressor proteins at relatively low levels applies also to the direct targets of EZH2. YAP and TAZ are extensively studied for their role as transcriptional co-activators, while their role as transcriptional co-repressors is still emerging but has not been fully clarified (32,43-45). Our findings help to dissect mechanistically the YAP/TAZ-mediated transcriptional repression of a subset of tumor-suppressor genes through the cooperation with EZH2. This was previously shown for the GDF15 gene in breast cancer cell lines (17), but we extend these findings also to other genes in NSCLC. The subset of common YAP/TAZ/PRC2 targets might be different with regard to cell and tissue contexts. Moreover, we did not find any physical interaction between YAP and EZH2 in our experimental model (data not shown). Future work will be aimed at looking for potential partners or protein complexes that may help the recruitment of YAP and EZH2 on the same genomic targets and mediate the formation of a repressive chromatin conformation. The NuRD complex due to its previously reported recruitment onto some YAP/TAZ repressed targets might be a strong candidate (32,44,45). The H3K27 de-acetylation mediated by the NuRD complex was shown in turn to recruit the PRC2 complex onto bivalent genes, in which both acetylation (associated with transcriptional activation) and methylation (associated with repression) of H3K27 are present. The balance between the two modifications determines the transcriptional state of the target genes (46). In agreement with this hypothesis, the depletion of YAP and TAZ reduces the enrichment of H3K27me3 and increases H3K27Ac at the YAP/TAZ/EZH2 co-repressed loci suggesting that YAP and TAZ may facilitate EZH2 mediated H3K27 methylation.

We also confirmed the transcriptional and functional synergism between YAP/TAZ and EZH2 in NSCLC through their combined inhibition upon pharmacological treatment with Dasatinib and Tazemetostat (EPZ-6438). Both are FDA approved drugs targeting YAP and EZH2 activity, respectively (34) (35). Tazemetostat has been tested in early phase trials for rhabdoid tumors, B-Cell Non-Hodgkin lymphoma, and has been recently approved for the treatment of epithelioid sarcoma (47). These tumors are characterized by overexpression or hyperactivation of EZH2 (48-50). However, in other cancer types, Tazemetostat was not as effective as a single therapeutic agent, suggesting the need to test the drug in combination with other anticancer agents (49). In NSCLC, Tazemetostat is currently evaluated in a clinical trial that involves multiple immunotherapy-based treatment combinations, both as the first-line and the second-line therapy of patients with confirmed metastases (ClinicalTrials.gov Identifier: NCT03337698). Dasatinib has failed in NSCLC clinical trials as single-agent due to its relatively high toxicity and it is currently used in the early phase trials in combination with EGFR inhibitors or immunotherapy (ClinicalTrials.gov Identifier: NCT02954523, NCT02750514). In our model (NSCLC cell lines), we observed a synergistic effect of Dasatinib and Tazemetostat treatment in the inhibition of cell proliferation and in the de-repression of tumor-suppression genes co-repressed by the combinatorial activity of YAP and EZH2.

## Conclusions

In sum, our findings suggest that the newly characterized YAP/TAZ/EZH2 oncogenic axis may represent a potential therapeutic target in lung cancer. More precisely, our results provide a rationale for a two-pronged strategy for inhibiting both YAP/TAZ as well as EZH2 for an efficient therapeutic outcome, because of their functional synergy.

## Acknowledgements

We thank Dr. Daniela Palacios (Santa Lucia Foundation, Rome Italy) for kindly providing the pCDNA3-EZH2 construct. We thank Dr. Stefen Wiemann (German Cancer Research Center (DKFZ), Heidelberg, Germany) for kindly sharing the plasmid for TGFBR2 3’UTR luciferase assay (Psi-check2-TGFBR2).

## References

1. Siegel RL, Miller KD, Jemal A. Cancer Statistics, 2017. CA: a cancer journal for clinicians 2017;67:7–30

2. Boloker G, Wang C, Zhang J. Updated statistics of lung and bronchus cancer in United States (2018). J Thorac Dis 2018;10:1158–61

3. Nadal E, Massuti B, Domine M, Garcia-Campelo R, Cobo M, Felip E. Immunotherapy with checkpoint inhibitors in non-small cell lung cancer: insights from long-term survivors. Cancer Immunol Immunother 2019

4. Camidge DR, Doebele RC, Kerr KM. Comparing and contrasting predictive biomarkers for immunotherapy and targeted therapy of NSCLC. Nat Rev Clin Oncol 2019

5. Zanconato F, Cordenonsi M, Piccolo S. YAP/TAZ at the Roots of Cancer. Cancer Cell 2016;29:783–803

6. Lo Sardo F, Forcato M, Sacconi A, Capaci V, Zanconato F, di Agostino S, et al. MCM7 and its hosted miR-25, 93 and 106b cluster elicit YAP/TAZ oncogenic activity in lung cancer. Carcinogenesis 2016

7. David CJ, Massague J. Contextual determinants of TGFbeta action in development, immunity and cancer. Nat Rev Mol Cell Biol 2018;19:419–35

8. Hata A, Chen YG. TGF-beta Signaling from Receptors to Smads. Cold Spring Harb Perspect Biol 2016;8

9. Lebrun JJ. The Dual Role of TGFbeta in Human Cancer: From Tumor Suppression to Cancer Metastasis. ISRN Mol Biol 2012;2012:381428

10. Gyorffy B, Surowiak P, Budczies J, Lanczky A. Online survival analysis software to assess the prognostic value of biomarkers using transcriptomic data in non-small-cell lung cancer. PLoS One 2013;8:e82241

11. Feng XH, Lin X, Derynck R. Smad2, Smad3 and Smad4 cooperate with Sp1 to induce p15(Ink4B) transcription in response to TGF-beta. EMBO J 2000;19:5178–93

12. Pardali K, Kurisaki A, Moren A, ten Dijke P, Kardassis D, Moustakas A. Role of Smad proteins and transcription factor Sp1 in p21(Waf1/Cip1) regulation by transforming growth factor-beta. J Biol Chem 2000;275:29244–56

13. Iavarone A, Massague J. Repression of the CDK activator Cdc25A and cell-cycle arrest by cytokine TGF-beta in cells lacking the CDK inhibitor p15. Nature 1997;387:417–22

14. Murai F, Koinuma D, Shinozaki-Ushiku A, Fukayama M, Miyaozono K, Ehata S. EZH2 promotes progression of small cell lung cancer by suppressing the TGF-beta-Smad-ASCL1 pathway. Cell Discov 2015;1:15026

15. Zhang H, Qi J, Reyes JM, Li L, Rao PK, Li F, et al. Oncogenic Deregulation of EZH2 as an Opportunity for Targeted Therapy in Lung Cancer. Cancer Discov 2016;6:1006–21

16. Kim MH, Kim J, Hong H, Lee SH, Lee JK, Jung E, et al. Actin remodeling confers BRAF inhibitor resistance to melanoma cells through YAP/TAZ activation. EMBO J 2016;35:462–78

17. Wang T, Mao B, Cheng C, Zou Z, Gao J, Yang Y, et al. YAP promotes breast cancer metastasis by repressing growth differentiation factor-15. Biochim Biophys Acta Mol Basis Dis 2018;1864:1744–53

18. Mizuno T, Murakami H, Fujii M, Ishiguro F, Tanaka I, Kondo Y, et al. YAP induces malignant mesothelioma cell proliferation by upregulating transcription of cell cycle-promoting genes. Oncogene 2012;31:5117–22

19. Oku Y, Nishiya N, Tazawa T, Kobayashi T, Umezawa N, Sugawara Y, et al. Augmentation of the therapeutic efficacy of WEE1 kinase inhibitor AZD1775 by inhibiting the YAP-E2F1-DNA damage response pathway axis. FEBS Open Bio 2018;8:1001–12

20. Zanconato F, Forcato M, Battilana G, Azzolin L, Quaranta E, Bodega B, et al. Genome-wide association between YAP/TAZ/TEAD and AP-1 at enhancers drives oncogenic growth. Nat Cell Biol 2015;17:1218–27

21. Bracken AP, Pasini D, Capra M, Prosperini E, Colli E, Helin K. EZH2 is downstream of the pRB-E2F pathway, essential for proliferation and amplified in cancer. EMBO J 2003;22:5323–35

22. Liu H, Li W, Yu X, Gao F, Duan Z, Ma X, et al. EZH2-mediated Puma gene repression regulates non-small cell lung cancer cell proliferation and cisplatin-induced apoptosis. Oncotarget 2016;7:56338–54

23. Sasaki M, Yamaguchi J, Itatsu K, Ikeda H, Nakanuma Y. Over-expression of polycomb group protein EZH2 relates to decreased expression of p16 INK4a in cholangiocarcinogenesis in hepatolithiasis. J Pathol 2008;215:175–83

24. Chen Z, Chen X, Chen P, Yu S, Nie F, Lu B, et al. Long non-coding RNA SNHG20 promotes non-small cell lung cancer cell proliferation and migration by epigenetically silencing of P21 expression. Cell Death Dis 2017;8:e3092

25. Sun Z, He C, Xiao M, Wei B, Zhu Y, Zhang G, et al. LncRNA FOXC2 antisense transcript accelerates non-small-cell lung cancer tumorigenesis via silencing p15. Am J Transl Res 2019;11:4552–60

26. Chen X, Wang K. lncRNA ZEB2-AS1 Aggravates Progression of Non-Small Cell Lung Carcinoma via Suppressing PTEN Level. Med Sci Monit 2019;25:8363–70

27. Tumaneng K, Schlegelmilch K, Russell RC, Yimlamai D, Basnet H, Mahadevan N, et al. YAP mediates crosstalk between the Hippo and PI(3)K-TOR pathways by suppressing PTEN via miR-29. Nat Cell Biol 2012;14:1322–9

28. Ma C, Wu G, Zhu Q, Liu H, Yao Y, Yuan D, et al. Long intergenic noncoding RNA 00673 promotes non-small-cell lung cancer metastasis by binding with EZH2 and causing epigenetic silencing of HOXA5. Oncotarget 2017;8:32696–705

29. Zhou X, Zang X, Ponnusamy M, Masucci MV, Tolbert E, Gong R, et al. Enhancer of Zeste Homolog 2 Inhibition Attenuates Renal Fibrosis by Maintaining Smad7 and Phosphatase and Tensin Homolog Expression. J Am Soc Nephrol 2016;27:2092–108

30. Lange AW, Sridharan A, Xu Y, Stripp BR, Perl AK, Whitsett JA. Hippo/Yap signaling controls epithelial progenitor cell proliferation and differentiation in the embryonic and adult lung. J Mol Cell Biol 2015;7:35–47

31. Lo Sardo F, Forcato M, Sacconi A, Capaci V, Zanconato F, Di Agostino S, et al. MCM7 and its hosted miR-25, 93 and 106b cluster elicit YAP/TAZ oncogenic activity in lung cancer. Carcinogenesis 2017;38:64–75

32. Kim M, Kim T, Johnson RL, Lim DS. Transcriptional co-repressor function of the hippo pathway transducers YAP and TAZ. Cell Rep 2015;11:270–82

33. Qin Z, Xia W, Fisher GJ, Voorhees JJ, Quan T. YAP/TAZ regulates TGF-beta/Smad3 signaling by induction of Smad7 via AP-1 in human skin dermal fibroblasts. Cell Commun Signal 2018;16:18

34. Oku Y, Nishiya N, Shito T, Yamamoto R, Yamamoto Y, Oyama C, et al. Small molecules inhibiting the nuclear localization of YAP/TAZ for chemotherapeutics and chemosensitizers against breast cancers. FEBS Open Bio 2015;5:542–9

35. Knutson SK, Wigle TJ, Warholic NM, Sneeringer CJ, Allain CJ, Klaus CR, et al. A selective inhibitor of EZH2 blocks H3K27 methylation and kills mutant lymphoma cells. Nat Chem Biol 2012;8:890–6

36. Zhao B, Ye X, Yu J, Li L, Li W, Li S, et al. TEAD mediates YAP-dependent gene induction and growth control. Genes Dev 2008;22:1962–71

37. Morera L, Lubbert M, Jung M. Targeting histone methyltransferases and demethylases in clinical trials for cancer therapy. Clin Epigenetics 2016;8:57

38. Sun J, Wang X, Tang B, Liu H, Zhang M, Wang Y, et al. A tightly controlled Src-YAP signaling axis determines therapeutic response to dasatinib in renal cell carcinoma. Theranostics 2018;8:3256–67

39. Wang M, Yuang-Chi Chang A. Molecular mechanism of action and potential biomarkers of growth inhibition of synergistic combination of afatinib and dasatinib against gefitinib-resistant non-small cell lung cancer cells. Oncotarget 2018;9:16533–46

40. Saito A, Nagase T. Hippo and TGF-beta interplay in the lung field. Am J Physiol Lung Cell Mol Physiol 2015;309:L756–67

41. Liu Y, He K, Hu Y, Guo X, Wang D, Shi W, et al. YAP modulates TGF-beta1-induced simultaneous apoptosis and EMT through upregulation of the EGF receptor. Sci Rep 2017;7:45523

42. Hiemer SE, Szymaniak AD, Varelas X. The Transcriptional Regulators TAZ and YAP Direct Transforming Growth Factor beta-induced Tumorigenic Phenotypes in Breast Cancer Cells. J Biol Chem 2014;289:13461–74

43. Hong JH, Hwang ES, McManus MT, Amsterdam A, Tian Y, Kalmukova R, et al. TAZ, a transcriptional modulator of mesenchymal stem cell differentiation. Science 2005;309:1074–8

44. Beyer TA, Weiss A, Khomchuk Y, Huang K, Ogunjimi AA, Varelas X, et al. Switch enhancers interpret TGF-beta and Hippo signaling to control cell fate in human embryonic stem cells. Cell Rep 2013;5:1611–24

45. Tan BS, Yang MC, Singh S, Chou YC, Chen HY, Wang MY, et al. LncRNA NORAD is repressed by the YAP pathway and suppresses lung and breast cancer metastasis by sequestering S100P. Oncogene 2019;38:5612–26

46. Reynolds N, Salmon-Divon M, Dvinge H, Hynes-Allen A, Balasooriya G, Leaford D, et al. NuRD-mediated deacetylation of H3K27 facilitates recruitment of Polycomb Repressive Complex 2 to direct gene repression. EMBO J 2012;31:593–605

47. First EZH2 Inhibitor Approved-for Rare Sarcoma. Cancer Discov 2020;10:333–4

48. Italiano A, Soria JC, Toulmonde M, Michot JM, Lucchesi C, Varga A, et al. Tazemetostat, an EZH2 inhibitor, in relapsed or refractory B-cell non-Hodgkin lymphoma and advanced solid tumours: a first-in-human, open-label, phase 1 study. Lancet Oncol 2018;19:649–59

49. Kurmasheva RT, Sammons M, Favours E, Wu J, Kurmashev D, Cosmopoulos K, et al. Initial testing (stage 1) of tazemetostat (EPZ-6438), a novel EZH2 inhibitor, by the Pediatric Preclinical Testing Program. Pediatr Blood Cancer 2017;64

50. Positive Results for Tazemetostat in Follicular Lymphoma. Cancer Discov 2018;8:OF3

