## Supplementary Fig. S1 for "YAP/TAZ and EZH2 synergize to impair tumor suppressor activity of TGFBR2 in non-small cell lung cancer"

### Figure S1

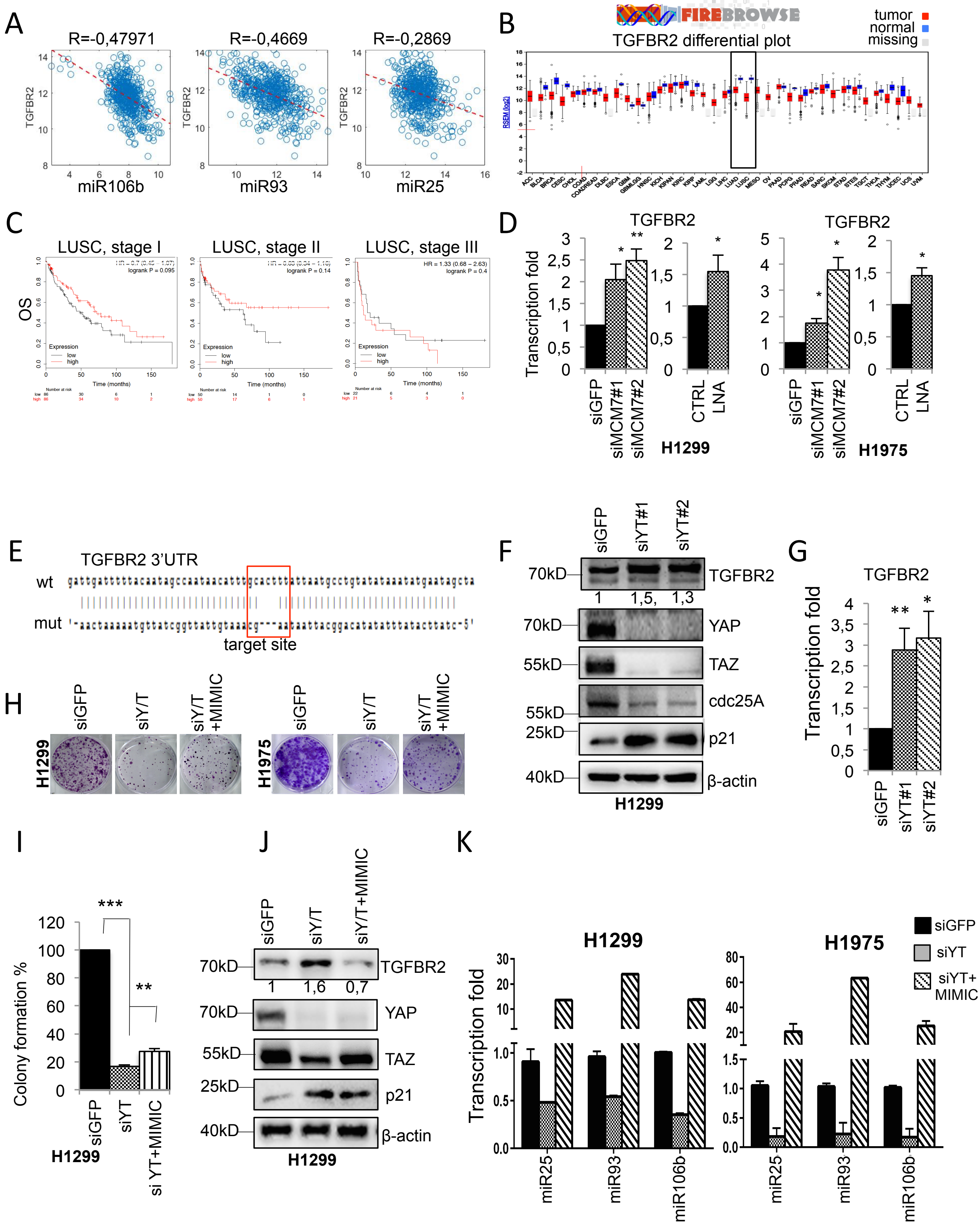

A
R=-0,47971
R=-0,4669
R=-0,2869
TGFBR2
miR106b
miR93
miR25
B
FIREBROWSE
TGFBR2 differential plot
tumor
normal
missing
C
LUSC, stage I
LUSC, stage II
LUSC, stage III
OS
Expression
low
high
Time (months)
D
TGFBR2
Transcription fold
siGFP
siMCM7#1
siMCM7#2
CTRL
LNA
H1299
H1975
E
TGFBR2 3'UTR
wt
mut
target site
F
siGFP
siYT#1
siYT#2
70kD
TGFBR2
1
1,5,
1,3
YAP
55kD
TAZ
cdc25A
25kD
p21
40kD
β-actin
H1299
G
TGFBR2
Transcription fold
siGFP
siYT#1
siYT#2
H
siGFP
siY/T
siY/T +MIMIC
H1299
H1975
I
Colony formation %
siGFP
siYT
si YT+MIMIC
H1299
J
siGFP
siY/T
siY/T+MIMIC
70kD
TGFBR2
1
1,6
0,7
YAP
55kD
TAZ
25kD
p21
40kD
β-actin
H1299
K
H1299
H1975
Transcription fold
siGFP
siYT
siYT+ MIMIC
miR25
miR93
miR106b

**Figure S1: TGFBR2 is a direct target of the oncogenic miR-106b-25 cluster and exhibits prognostic value in NSCLC.**

**A,** Dot plot showing the correlation between TGFBR2 transcript and miR-25, 93, and 106b in LUng Adenocarcinoma patients from the TCGA. **B,** Boxplots of TGFBR2 expression profile in different cancer types from patients samples deposited in the TCGA casuistry as obtained from the FireBrowse Gene Expression Viewer (http//firebrowse.org). Lung Adenocarcinoma (LUAD) tumor samples: 517, first quartile: 11,2, Median: 11,8, Third quartile: 12,3, Fold change: 0,287. Lung Squamous Cell Carcinoma (LUSC) tumor samples: 501, First quartile: 9,79, Median: 10,5, third quartile: 11,2, fold change: 0,117. **C,** KM of lung squamous cell carcinoma (LUSC) patients stratified for tumor stage, with high or low expression of TGFBR2. The number of patients is indicated below the plots. **D,** Real-time PCR analysis of TGFBR2 transcript in H1299 cells (left) and H1975 cells (right) upon the interference of MCM7 or transfection with LNA miR-25, 93, and 106b. GAPDH transcript was used for normalization. Each experiment was performed at least in triplicate. Two-tailed t-test analysis was applied to calculate the P values. *p<0,05; **p<0,01;***p<0,001. **E,** Schematic representation of the 3' UTR of TGFBR2 transcript with the sequence targeted by miR-93 and miR-106b in the wt and mutant version. **F-G,** Western blot analysis of the indicated proteins (**F**) and real-time PCR analysis of TGFBR2 transcript, normalized to GAPDH (**G**) in H1299 cells upon interference with alternative siRNAs against YAP and TAZ. Numbers below TGFBR2 blot represent the relative abundance of TGFBR2 protein normalized to β-actin as measured by densitometry (Uvitec) and normalized to B-actin signal. **H,** Representative image of cell colony formation assay in H1299 (left panels) and H1975 (right panels) upon YAP/TAZ interference with or without overexpression of mimic miR-25, 93, and 106b. **I,** quantification of colonies in H1299 cells upon YAP/TAZ interference with or without overexpression of mimic miR-25, 93, and 106b. **J,** Western blot of the indicated proteins in H1299 cells upon YAP/TAZ interference with or without overexpression of mimic miR-25, 93, and 106b. **K,** Taq-Man based quantification of miR-25, 93, and 106b, normalized to RNU48 and RNU49, in H1299 (left) and H1975 (right) upon YAP/TAZ interference with or without overexpression of mimic miR-25, 93 and 106b.
