## Supplementary Fig. S2 for "YAP/TAZ and EZH2 synergize to impair tumor suppressor activity of TGFBR2 in non-small cell lung cancer"

### Figure S2

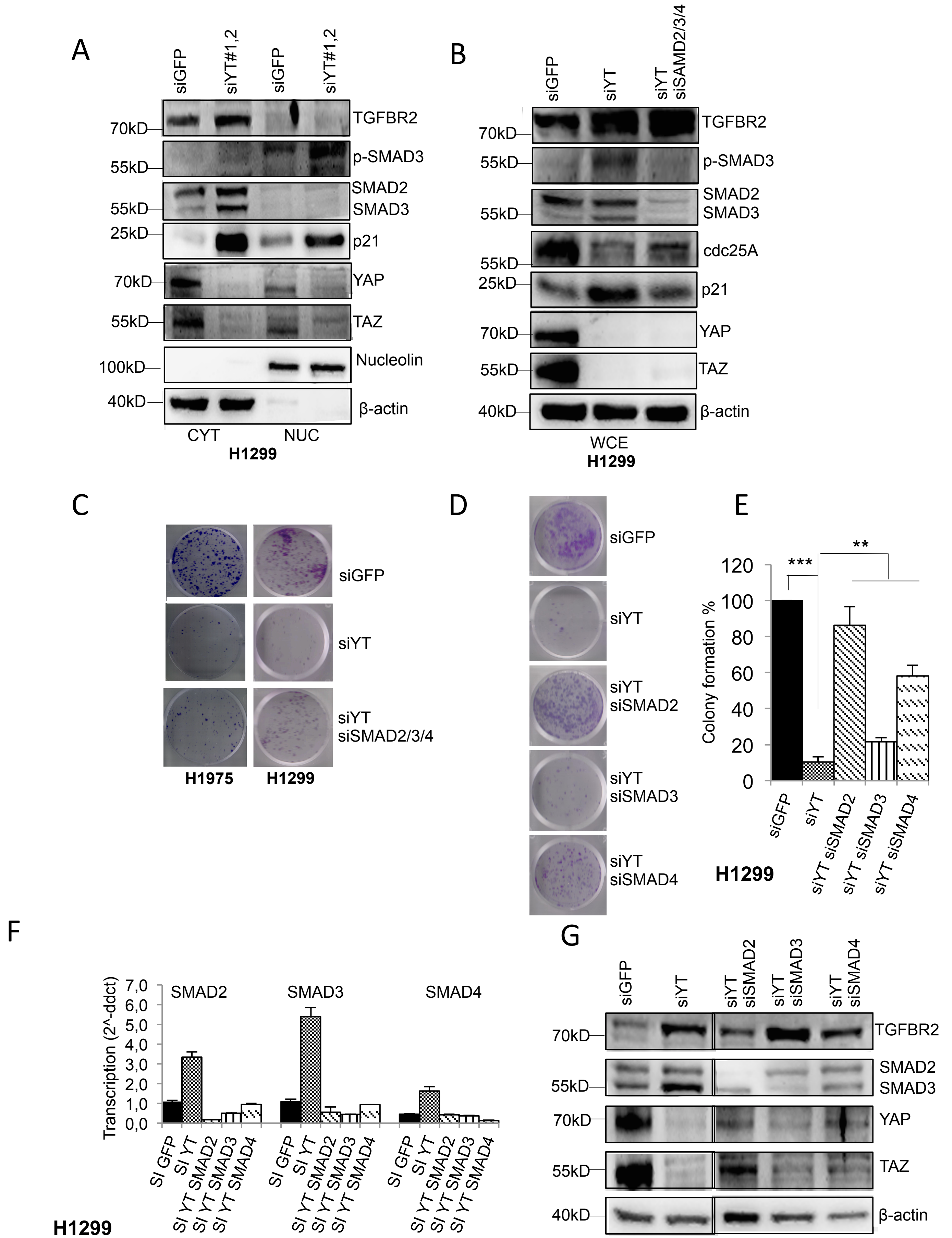

A
siGFP
siYT#1,2
siGFP
siYT#1,2
TGFBR2
p-SMAD3
SMAD2
SMAD3
p21
YAP
TAZ
Nucleolin
β-actin
70kD
55kD
55kD
25kD
70kD
55kD
100kD
40kD
CYT
NUC
H1299
B
siGFP
siYT
siYT siSAMD2/3/4
TGFBR2
p-SMAD3
SMAD2
SMAD3
cdc25A
p21
YAP
TAZ
β-actin
WCE
H1299
C
siGFP
siYT
siYT siSMAD2/3/4
H1975
H1299
D
siGFP
siYT
siYT siSMAD2
siYT siSMAD3
siYT siSMAD4
E
***
**
Colony formation %
siGFP
siYT
siYT siSMAD2
siYT siSMAD3
siYT siSMAD4
H1299
F
Transcription (2^-ddct)
SMAD2
SMAD3
SMAD4
SI GFP
SI YT
SI YT SMAD2
SI YT SMAD3
SI YT SMAD4
H1299
G
siGFP
siYT
siYT siSMAD2
siYT siSMAD3
siYT siSMAD4
TGFBR2
SMAD2
SMAD3
YAP
TAZ
β-actin

**Figure S2: YAP/TAZ control TGFBR2 expression acting upstream to the miR cluster.**

**A,** Western blot analysis of nucleo-cytoplasmic extracts from H1299 cells showing the abundance of the indicated proteins upon YAP/TAZ interference compared to siGFP control cells. Nucleolin and β-actin were used as a nuclear and cytoplasmic loading control, respectively. **B,** Western blot analysis of whole-cell extracts (WCE) of the indicated proteins normalized to B-actin in H1299 cells upon the interference of YAP and TAZ with two different combinations of alternative siRNAs with or without concomitant interference of SMADs. **C,** Representative image of colony formation assay in H1975 (left) and H1299 cells (right) upon YAP/TAZ interference with or without concomitant interference of SMAD2/3/4. **D-E,** Representative image (**D**) and relative quantification chart (**E**) of colony formation in H1299 cells upon YAP/TAZ interference with or without concomitant interference of SMAD2, SMAD3 or SMAD4, respect to siGFP control cells. **F,** Real-time quantification of SMAD2, SMAD3 and SMAD4 transcripts, normalized to GAPDH, in H1299 upon the interference of YAP/TAZ with or without concomitant interference of SMAD2, SMAD3 and SMAD4. **G,** Western blot analysis of the indicated proteins, normalized to β-actin, in H1299 upon the interference of YAP/TAZ with or without concomitant interference of SMAD2, SMAD3 and SMAD4.
