## Supplementary Fig. S3 for "YAP/TAZ and EZH2 synergize to impair tumor suppressor activity of TGFBR2 in non-small cell lung cancer"

**Figure S3**

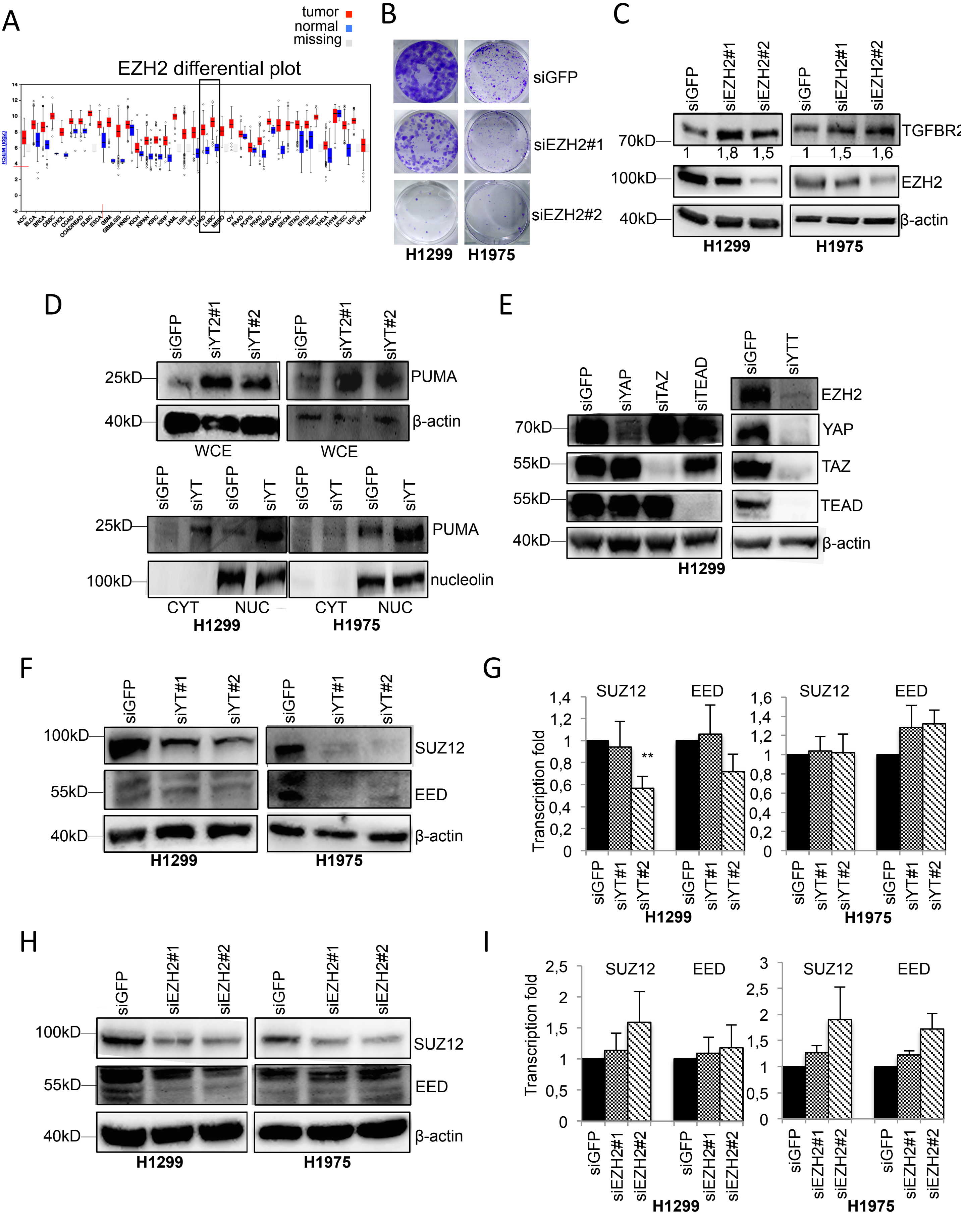

A
tumor
normal
missing
EZH2 differential plot
B
siGFP
siEZH2#1
siEZH2#2
H1299
H1975
C
siGFP
siEZH2#1
siEZH2#2
70kD
100kD
40kD
TGFBR2
EZH2
β-actin
1
1,8
1,5
1
1,5
1,6
H1299
H1975
D
siGFP
siYT2#1
siYT#2
25kD
40kD
PUMA
β-actin
WCE
siYT
100kD
nucleolin
CYT
NUC
H1299
H1975
E
siYAP
siTAZ
siTEAD
siYTT
55kD
YAP
TAZ
TEAD
F
siYT#1
SUZ12
EED
G
Transcription fold
SUZ12
EED
**
H
I

**Figure S3: EZH2 is oncogenic in NSCLC and is a transcriptional target of YAP/TAZ/TEAD. A,** Boxplots of EZH2 expression profile in different cancer types from patient samples deposited in the TCGA casuistry as obtained from the FireBrowse Gene Expression Viewer. **B,** Representative images of colony formation of H1299 (left) and H1975 (right) upon EZH2 interference with alternative siRNAs. **C,** Western blot analysis of the indicated proteins and quantification by densitometry normalized to β-actin in H1299 (left) and H1975 (right) upon EZH2 interference with alternative siRNAs. **D,** Western blot analysis of Whole Cell Extract (WCE, upper panel) and nucleo-cytoplasmic extract (lower panel) of the indicated proteins in H1299 cells (left) and H1975 (right) upon YAP/TAZ interference. β-actin and nucleolin were used as a loading control, respectively, for WCE and nucleo-cytoplasmic extract. **E,** Western blot analysis of the indicated proteins in H1299 cells upon YAP, TAZ and TEAD interference. (Interference control of a representative ChIp experiment). **F,** Western blot analysis of the indicated proteins, normalized to β-actin, in H1299 (left panels) and H1975 (right panels) upon YAP/TAZ interference with two different combinations of alternative siRNAs. **G,** Real-time PCR quantification of the indicated transcripts, normalized to GAPDH, in H1299 (left panels) and H1975 (right panels) upon YAP/TAZ interference. **H,** Western blot analysis of the indicated proteins, normalized to β-actin, in H1299 (left panels) and H1975 (right panels) upon EZH2 interference with two alternative siRNA. **I,** Real-time PCR quantification of the indicated transcripts, normalized to GAPDH, in H1299 (left panels) and H1975 (right panels) upon EZH2 interference.
