## Supplementary Fig. S4 for "YAP/TAZ and EZH2 synergize to impair tumor suppressor activity of TGFBR2 in non-small cell lung cancer"

### Figure S4

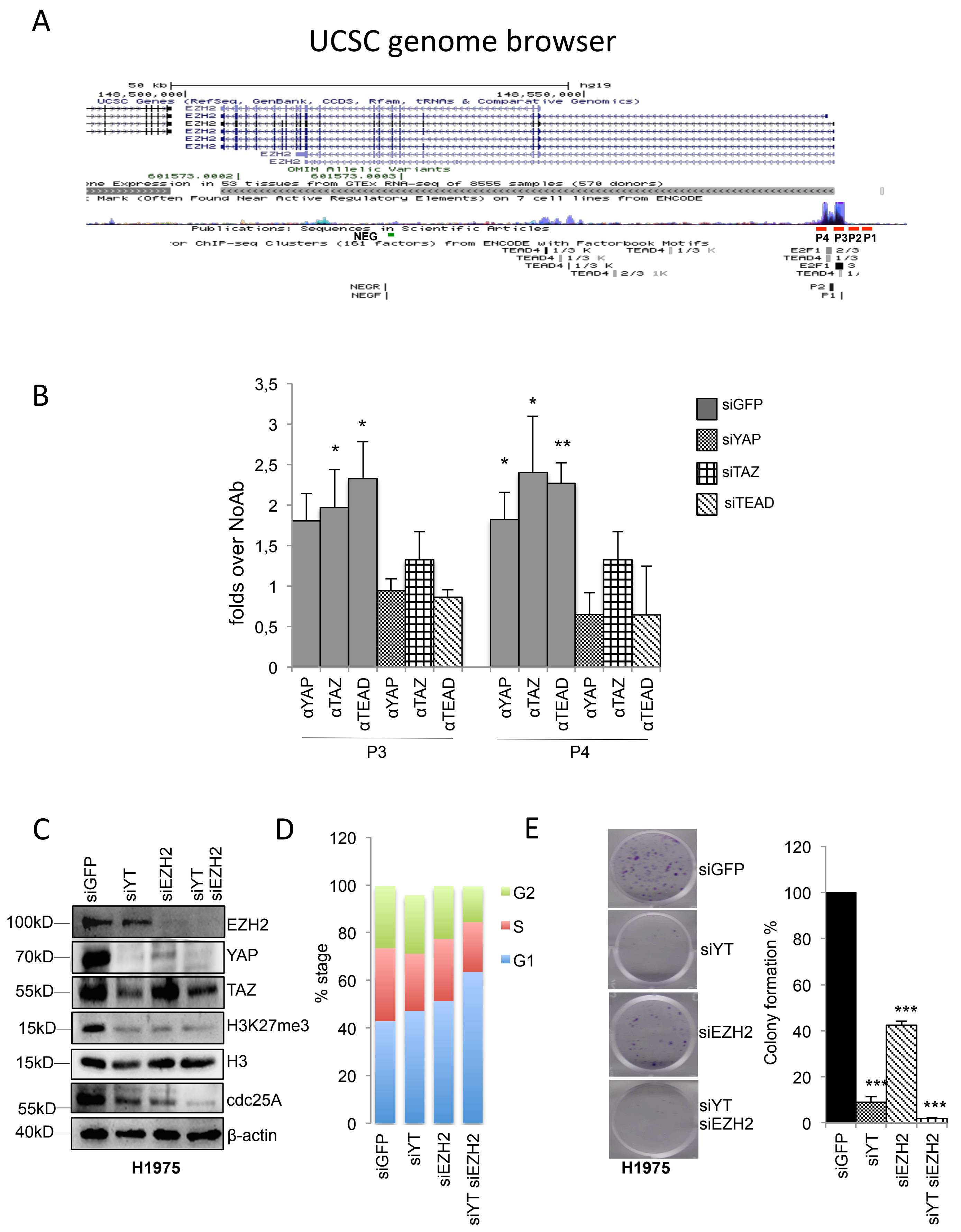

A
UCSC genome browser
NEG
P4 P3 P2 P1
B
3,5
3
2,5
2
1,5
1
0,5
0
folds over NoAb
*
*
*
*
**
siGFP
siYAP
siTAZ
siTEAD
αYAP
αTAZ
αTEAD
αYAP
αTAZ
αTEAD
P3
αYAP
αTAZ
αTEAD
αYAP
αTAZ
αTEAD
P4
C
siGFP
siYT
siEZH2
siYT siEZH2
100kD
70kD
55kD
15kD
15kD
55kD
40kD
EZH2
YAP
TAZ
H3K27me3
H3
cdc25A
β-actin
H1975
D
120
100
80
60
40
20
0
% stage
G2
S
G1
siGFP
siYT
siEZH2
siYT siEZH2
E
siGFP
siYT
siEZH2
siYT siEZH2
H1975
120
100
80
60
40
20
0
Colony formation %
***
***
***
siGFP
siYT
siEZH2
siYT siEZH2

**Figure S4: YAP/TAZ and EZH2 synergistically affect cell growth of lung cancer cell lines.**

**A,** Representation of the EZH2 locus from the UCSC Genome Browser, with the H7K27Ac, TEAD4, and E2F1 ChIp-Seq peaks from Encode at the EZH2 promoter. P1 and P2, P3 and P4 regions are marked with red lines, negative control region is marked with a green line. **B,** ChIP Fold enrichment of YAP, TAZ, and TEAD1 proteins onto the indicated sites of EZH2 locus in H1299 cells depleted for YAP, TAZ, and TEAD1 compared to control cells. Data are presented as mean ± SEM of at least three biological replicates. For each antibody, fold enrichment was calculated over no antibody control. **C,** Western blot analysis of the indicated proteins in H1975 cells upon the interference of YAP/TAZ and EZH2, either alone or in combination, respect to control cells. **D,** Cells (%) in G1, S and G2 phases in H1975 upon the interference of YAP/TAZ and EZH2, either alone or in combination, respect to control cells. **E,** Representative image (left) and quantification (right) of colony formation in H1975 cells, upon the interference of YAP/TAZ or EZH2, alone or in combination, respect to control cells.
