## Supplementary Fig. S5 for "YAP/TAZ and EZH2 synergize to impair tumor suppressor activity of TGFBR2 in non-small cell lung cancer"

Figure S5.

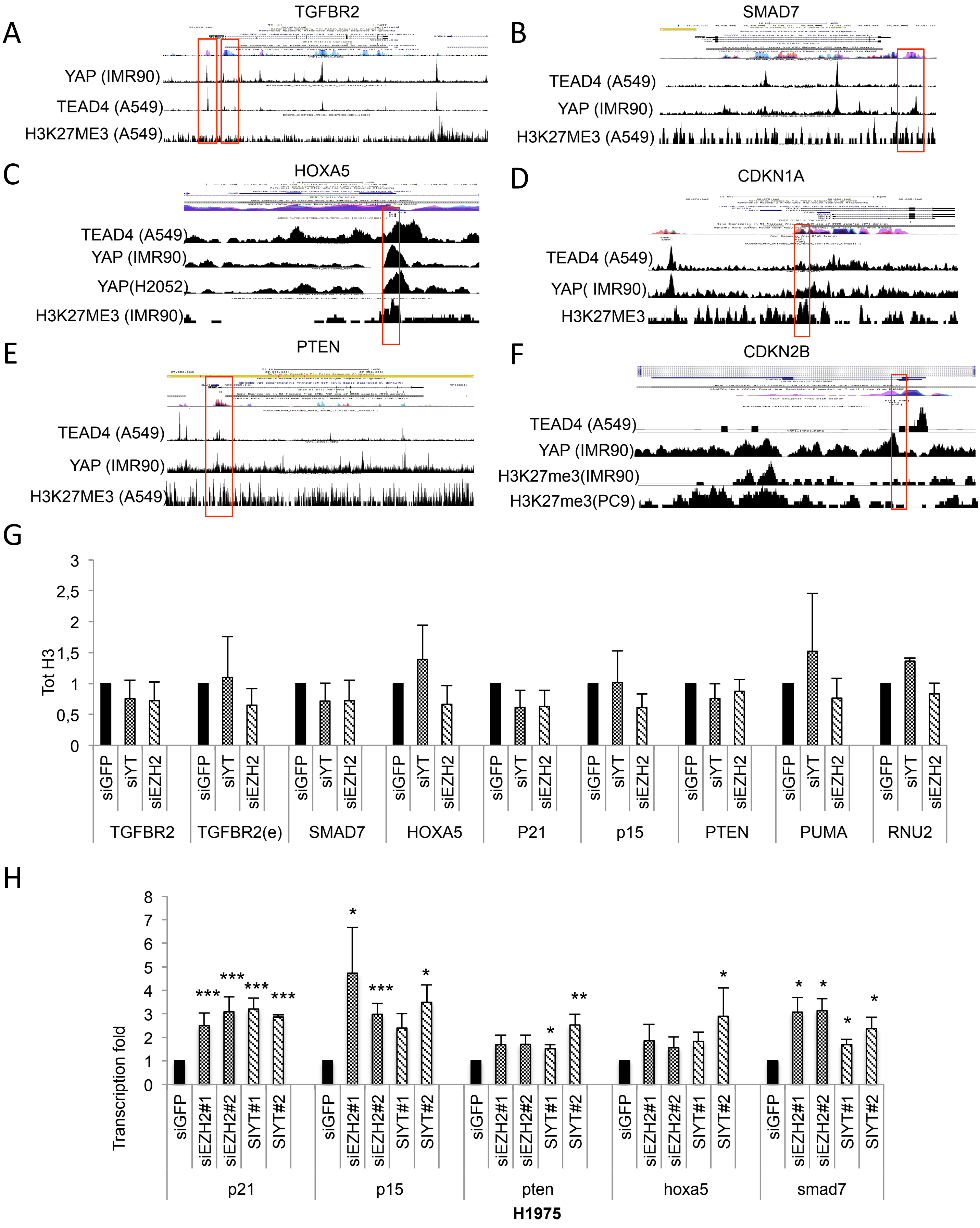

A
TGFBR2
YAP (IMR90)
TEAD4 (A549)
H3K27ME3 (A549)
B
SMAD7
TEAD4 (A549)
YAP (IMR90)
H3K27ME3 (A549)
C
HOXA5
TEAD4 (A549)
YAP (IMR90)
YAP(H2052)
H3K27ME3 (IMR90)
D
CDKN1A
TEAD4 (A549)
YAP( IMR90)
H3K27ME3
E
PTEN
TEAD4 (A549)
YAP (IMR90)
H3K27ME3 (A549)
F
CDKN2B
TEAD4 (A549)
YAP (IMR90)
H3K27me3(IMR90)
H3K27me3(PC9)
G
Tot H3
3
2,5
2
1,5
1
0,5
0
siGFP
siYT
siEZH2
TGFBR2
TGFBR2(e)
SMAD7
HOXA5
P21
p15
PTEN
PUMA
RNU2
H
Transcription fold
8
7
6
5
4
3
2
1
0
siGFP
siEZH2#1
siEZH2#2
SIYT#1
SIYT#2
p21
p15
pten
hoxa5
smad7
H1975

**Figure S5: Some onco-suppressive genes are co-repressed and co-occupied by YAP/TAZ/ EZH2 in NSCLC.**

**A-F,** UCSC Genome Browser tracks of YAP, TEAD4, and H3K27me3 in the indicated cell lines on the indicated genes as obtained through the Cistrome Browser DB. Red boxes indicate regions amplified for ChIp analysis. For TGFBR2 locus, both the promoter and the enhancer (e) were analyzed. **G,** ChIp fold enrichment of total H3 on the indicated loci. TGFBR2(e) indicates the enhancer region shown in figure S5a. Fold enrichment was calculated over no antibody control and normalized to the siGFP control that was adjusted to 1. Experiments were performed in triplicate. **H,** Real-time PCR analysis of the indicated transcripts in H1975 cells upon depletion of either YAP/ TAZ or EZH2 proteins.
