## Supplementary Fig. S6 for "YAP/TAZ and EZH2 synergize to impair tumor suppressor activity of TGFBR2 in non-small cell lung cancer"

**Figure S6**

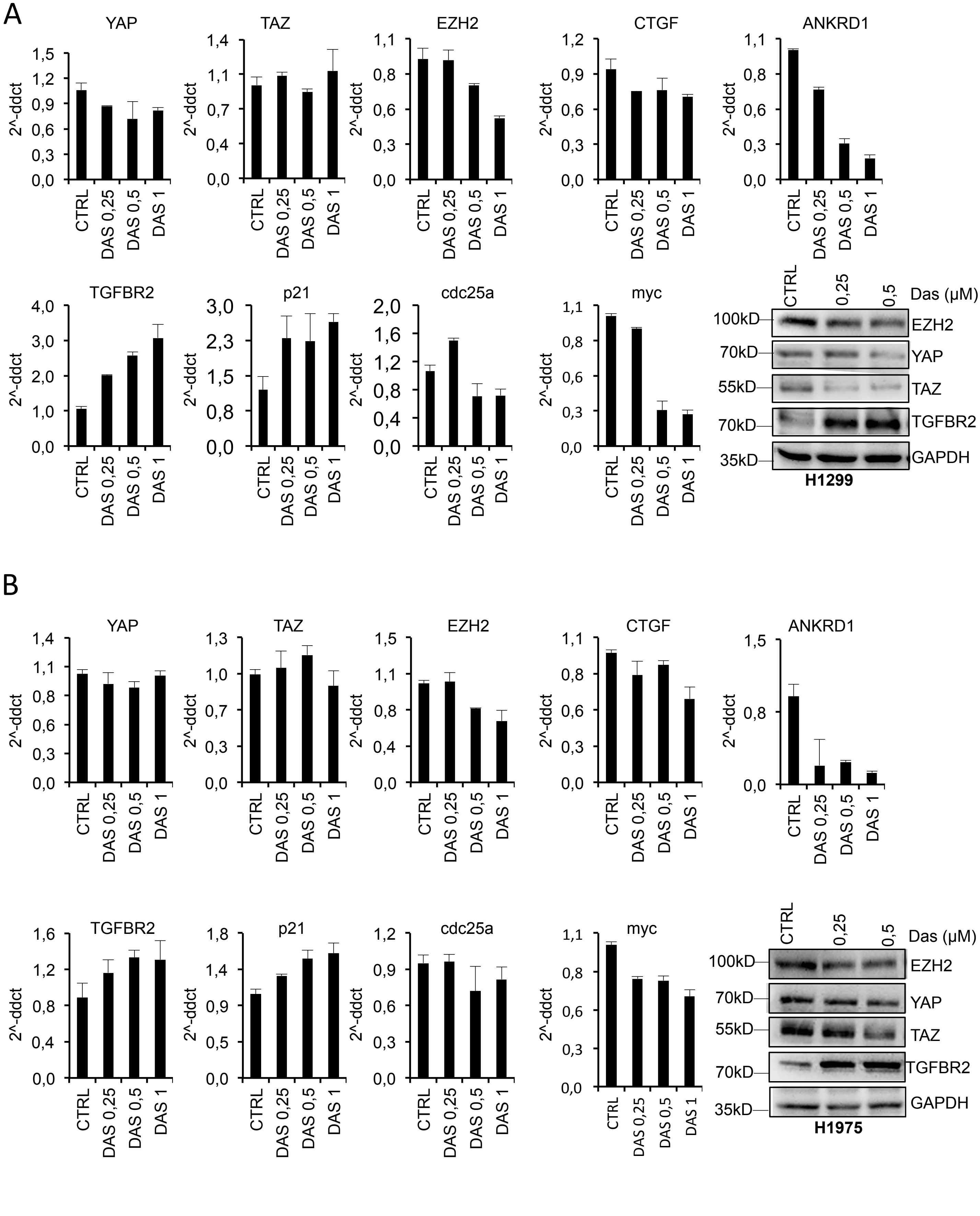
A
YAP
TAZ
EZH2
CTGF
ANKRD1
TGFBR2
p21
cdc25a
myc
2^-ddct
CTRL
DAS 0,25
DAS 0,5
DAS 1
CTRL
0,25
0,5
Das (µM)
100kD
70kD
55kD
70kD
35kD
EZH2
YAP
TAZ
TGFBR2
GAPDH
H1299
B
YAP
TAZ
EZH2
CTGF
ANKRD1
TGFBR2
p21
cdc25a
myc
2^-ddct
CTRL
DAS 0,25
DAS 0,5
DAS 1
CTRL
0,25
0,5
Das (µM)
100kD
70kD
55kD
70kD
35kD
EZH2
YAP
TAZ
TGFBR2
GAPDH
H1975

**Figure S6: Dasatinib treatment reduces YAP and TAZ transcriptional activity in NSCLC.**
**A-B,** Real-time PCR quantification of the indicated transcripts normalized to GAPDH (charts) and western blot analysis of the indicated proteins, normalized to β-actin (right panel) in H1299 (**A**) and H1975 cells (**B**) upon treatment with Dasatinib at the indicated doses. Charts are representative of one biological replicate and the experiments were repeated twice.
