## Supplementary Fig. S7 for "YAP/TAZ and EZH2 synergize to impair tumor suppressor activity of TGFBR2 in non-small cell lung cancer"

**Figure S7**

A

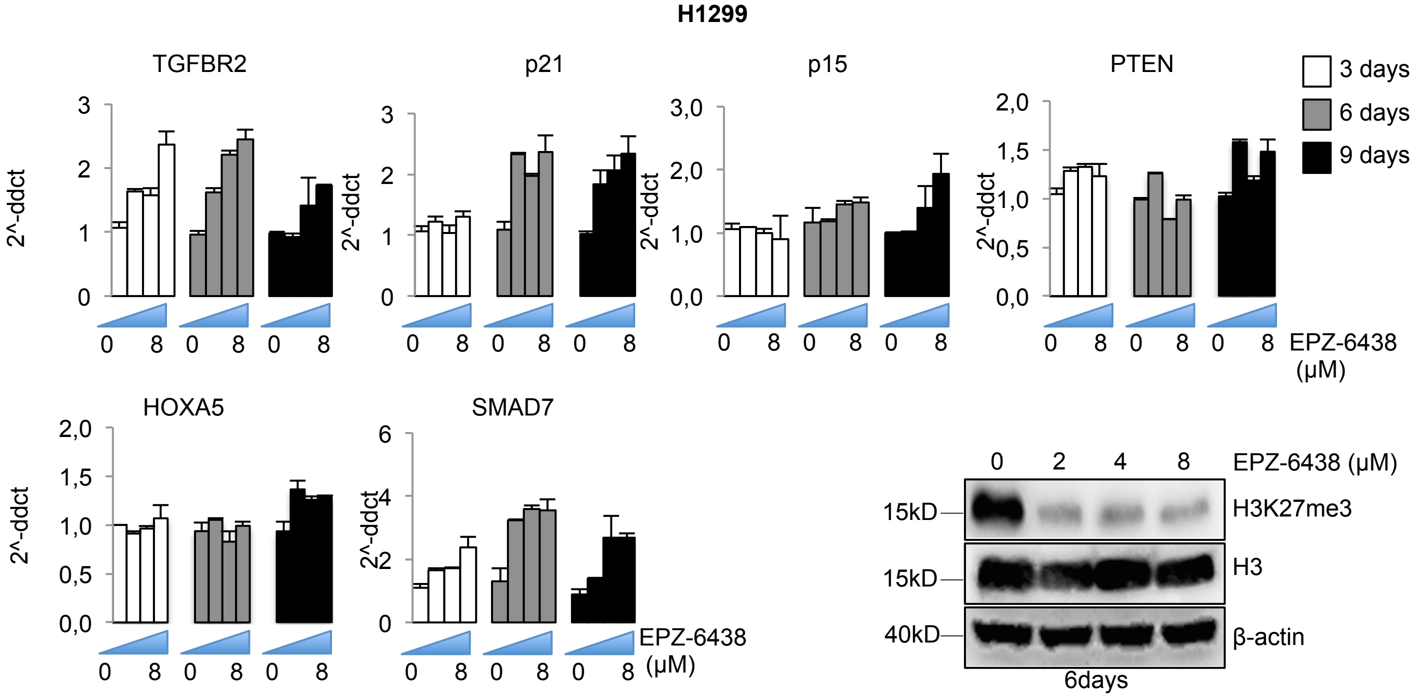
H1299
TGFBR2
p21
p15
PTEN
HOXA5
SMAD7
2^-ddct
3 days
6 days
9 days
0 8 0 8 0 8
EPZ-6438 (µM)
0 2 4 8
15kD
40kD
H3K27me3
H3
β-actin
6days

B

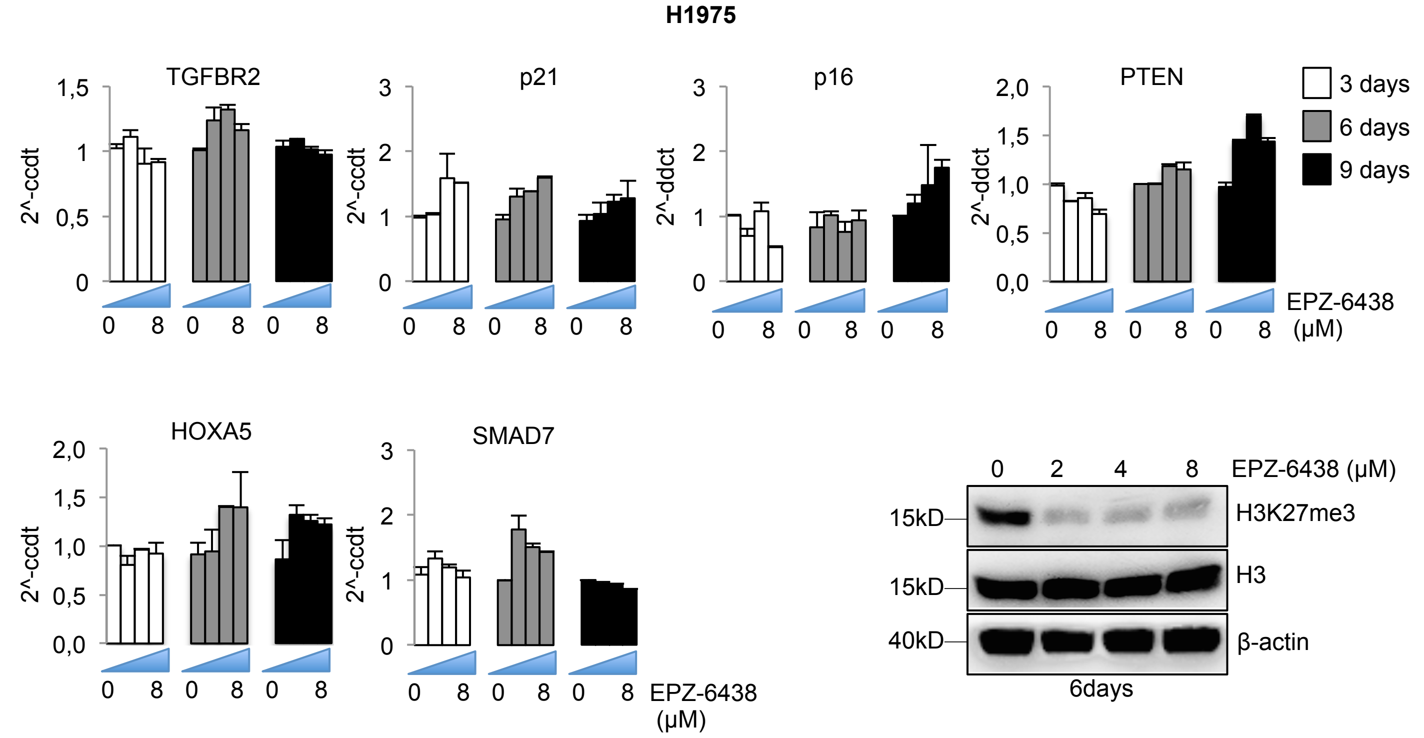
H1975
TGFBR2
p21
p16
PTEN
HOXA5
SMAD7
2^-ccdt
2^-ddct
3 days
6 days
9 days
0 8 0 8 0 8
EPZ-6438 (µM)
0 2 4 8
15kD
40kD
H3K27me3
H3
β-actin
6days

**Figure S7: Tazemetostat treatment de-represses PRC2 targets.**

**A-B,** Real-time PCR quantification of the indicated transcripts normalized to GAPDH and western blot analysis of the indicated proteins, normalized to β-actin (right panel) in H1299 (**A**) and H1975 cells (**B**) upon treatment with the EZH2 inhibitor Tazemetostat (EPZ-6438) at the indicated doses and time points.
