## Supplementary Fig. S8 for "YAP/TAZ and EZH2 synergize to impair tumor suppressor activity of TGFBR2 in non-small cell lung cancer"

Figure S8

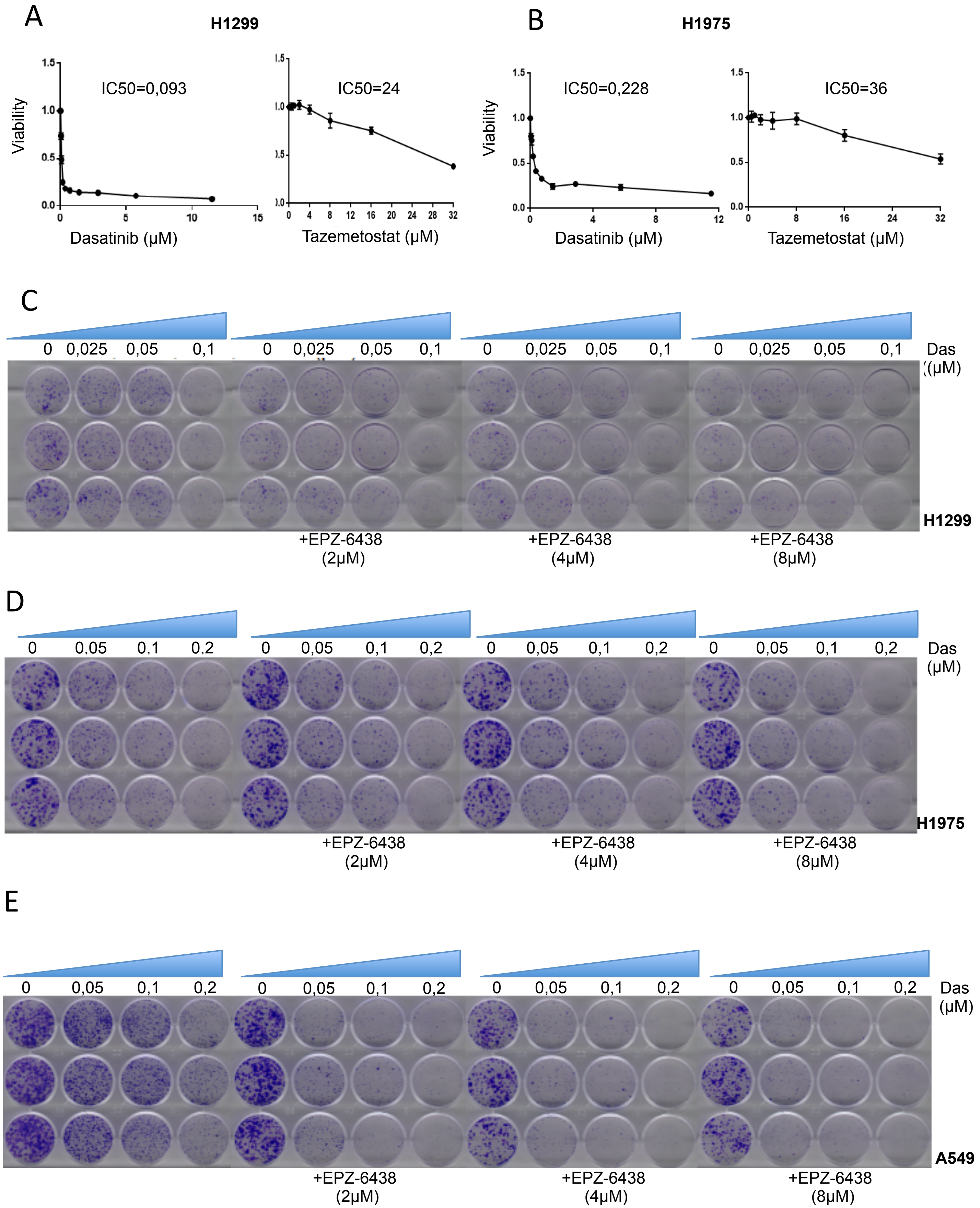
A
H1299
Viability
IC50=0,093
Dasatinib (µM)
IC50=24
Tazemetostat (µM)
B
H1975
Viability
IC50=0,228
Dasatinib (µM)
IC50=36
Tazemetostat (µM)
C
0 0,025 0,05 0,1
Das ((µM)
H1299
+EPZ-6438 (2µM)
+EPZ-6438 (4µM)
+EPZ-6438 (8µM)
D
0 0,05 0,1 0,2
Das (µM)
H1975
+EPZ-6438 (2µM)
+EPZ-6438 (4µM)
+EPZ-6438 (8µM)
E
0 0,05 0,1 0,2
Das (µM)
A549
+EPZ-6438 (2µM)
+EPZ-6438 (4µM)
+EPZ-6438 (8µM)

**Figure S8: Low doses of Dasatinib with Tazemetostat synergistically affect cell proliferation. A-B,** Viability of H1299 cells (**A**) and H1975 cells (**B**) as measured with ATPlite assay after 72h treatment with Dasatinib (left panel) or Tazemetostat (right panel) at the indicated doses. **C-E**, The complete panel of colony formation in H1299 (**C**) H1975 (**D**) and A549 cells (**E**) upon treatment with different combinations of Dasatinib and Tazemetostat at the indicated doses.
