## Supplementary Fig. S9 for "YAP/TAZ and EZH2 synergize to impair tumor suppressor activity of TGFBR2 in non-small cell lung cancer"

### Figure S9

A

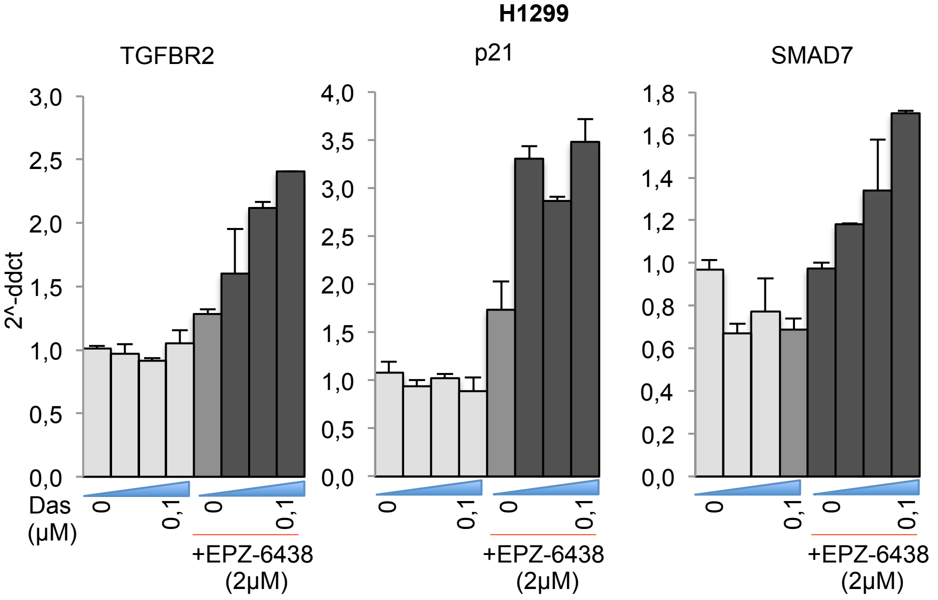
H1299
TGFBR2
p21
SMAD7
2^-ddct
3,0
2,5
2,0
1,5
1,0
0,5
0,0
4,0
3,5
3,0
2,5
2,0
1,5
1,0
0,5
0,0
1,8
1,6
1,4
1,2
1,0
0,8
0,6
0,4
0,2
0,0
Das (μM)
0
0,1
+EPZ-6438 (2μM)

B

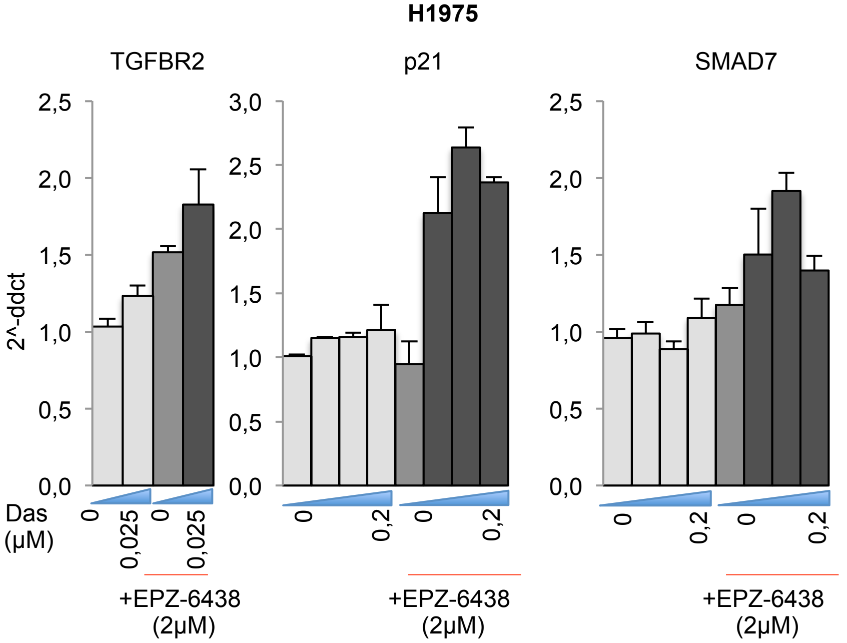
H1975
TGFBR2
p21
SMAD7
2^-ddct
2,5
2,0
1,5
1,0
0,5
0,0
3,0
2,5
2,0
1,5
1,0
0,5
0,0
Das (μM)
0
0,025
0,2
+EPZ-6438 (2μM)

**Figure S9: Dasatinib and Tazemetostat synergistically derepress oncosuppressor genes in NSCLS.**

**A-B,** Real-time PCR quantification of the indicated transcripts, normalized to GAPDH, in H1299 (**A**) and H1975 (**B**) after 72h treatment with increasing doses of Dasatinib with or without 2μM Tazemetostat.
