## Supplementary materials and Methods for "YAP/TAZ and EZH2 synergize to impair tumor suppressor activity of TGFBR2 in non-small cell lung cancer"

### Luciferase assay

For Luciferase assay, H1299 or H1975 cells were co-transfected in 24-well dishes using Lipofectamine 2000 (Invitrogen) with 100 ng of psiCheck2 containing a TGFBR2 3′ UTR-Renilla luciferase reporter, the same vector with a 3bp deletion in the seed sequence recognized by miR-93 and 106b or psiCheck2 empty vector as a control, plus the mirVana® *miRNA* mimic for miR-93 and miR-106b or a mimic negative control at a final concentration of 5nM. Cells were harvested 48h post transfection and luciferase activities were analyzed by the dual-luciferase reporter assay system (Promega, Madison, WI) in the GloMax 96 Microplate Luminometer (Promega). Mutagenesis was performed through QuikChange II Site-Directed Mutagenesis Kit (Agilent technologies) following the manufacturer's instructions and using the nucleotides listed below: TGFBR2 3UTR del94-96 F: 5'-ttgatttttacaatagccaataacatttgcttattaatgcctgtatataaatatgaatag-3' TGFBR2 3UTR del94-96 R: 5'-ctattcatatttatatacaggcattaataagcaaatgttattggctattgtaaaaatcaa-3' Correct deletion was confirmed by sequencing.

### Chromatin Immunoprecipitation (ChIp)

Cells were fixed with 1% formaldehyde (Sigma) in culture medium for 10 min at room temperature, and chromatin from lysed nuclei was sheared to 200–600 bp fragments using a Bioruptor sonicator (Diagenode). 100 μg of sheared chromatin and 5 μg of antibody plus 40 μl of magnetic beads (Dynabeads. Protein G 10004D ThermoFisher) were used for each Ip. Rabbit monoclonal anti YAP (Santa Cruz, sc-15407), anti TAZ (Sigma anti-WWTR1, HPA007415), anti TEAD (BD-Transduction Laboratories, 610923), anti-H4Ac, anti H3K27me3, Anti H3K27Ac, or no antibody as negative control were used. Quantitative real-time PCR was carried out with an Applied Biosystems® 7500 fast, *StepOne* Real-

Time PCR or QuantStudio 5 and 7 Real-Time PCR. Each sample was analyzed in triplicate. The amount of immunoprecipitated DNA in each sample was determined as the fraction of the input [amplification efficiency(Ct ChIP- Ct Input)], and normalized to the negative control (No Antibody).

**List of antibodies used for Western Blotting**

Western blotting was performed using the following primary antibodies: rabbit polyclonal anti YAP (Santa Cruz, sc-15407, 1:800), mouse monoclonal anti YAP 1A12 (Cell Signaling, 12395), rabbit monoclonal rabbit polyclonal anti TAZ (Sigma anti-WWTR1, HPA007415), mouse monoclonal anti B-actin (ACTBD11B7, Santa Cruz, sc-81178), mouse monoclonal anti c-Myc 9E10 (Santa Cruz, Sc-40) rabbit monoclonal anti MCM7 (D10A11, Cell Signaling, 3735S), mouse monoclonal anti-TEF-1 (BD-Transduction Laboratories, 610923), rabbit monoclonal anti-p21 Waf1/Cip1 (Cell Signaling, 2947), rabbit polyclonal anti-TGFBR2 (C-20, Santa-Cruz sc-220, 1:200), mouse monoclonal anti-cdc25A (DCS-120, Santa Cruz, sc-56264, 1:500), mouse monoclonal anti-p16 (F-12, Santa Cruz, sc-1661) rabbit polyclonal anti-smad2-3 (Cell Signaling, 3102), rabbit monoclonal anti-phospho smad3 (Ser423/425, C25A9 Cell Signaling, 9520) rabbit polyclonal anti-PUMA (NT) bbc3 (Assay Designs, Ann Arbor, 905-237), rabbit polyclonal anti EZH2 (Bethyl laboratories, A304-197A), rabbit polyclonal Anti SUZ12 (Bethyl laboratories, A302-407A), rabbit monoclonal Anti SUZ12 (Cell Signaling, 3737) rabbit monoclonal anti EED (Cell Signaling 51673), mouse monoclonal anti H3K27me3 (Abcam Ab 6002), rabbit polyclonal anti-acetyl H3K27 (Sigma-Aldrich, 07-360), rabbit polyclonal anti Acetil H4 (Sigma-Aldrich, 06-598). Where not specified differently, primary antibodies were diluted 1:1000 in TBS with 5%BSA and 0,1% tween. Secondary antibodies used were goat anti-mouse and goat anti-rabbit, conjugated to horseradish peroxidase (Amersham Biosciences, Piscataway, NJ) diluted 1:10000 in TBS with 5%BSA and 0,1%

tween. Precision Plus *Protein Standard was used to monitor the protein run and tranfer and to estimate* the protein Molecular weight.

**Sequence of siRNA used for interference**

siGFP (as non-silencing control) 5′-AAGUUCAGCGUGUCCGGGGAG-3′, siYAP#1: 5′ GACAUCUUCUGGUCAGAGA-3′, siYAP#2: 5′-CUGGUCAGAGAUACUUCUU-3′, siTAZ#1: 5′-AAAGUUCCUAAGUCAACGU-3′, siTAZ#2: 5′-AGGUACUUCCUCAAUCACA-3′, siMCM7#1: 5′-UACUACGAGGGAUAUUUCCUU-3′, siMCM7#2: 5″-GAUCACACGAGGCUUGUUGUU-3′, siTEAD1#1: 5′-CGAUUUGUAUACCGAAUAA-3′, siTEAD1#2: 5′-GAAAGGUGGCUUAAAGGAA-3′, siTGFBR2#1: 5’-GATTCAAGAGTATTCTCACTT-3’, siTGFBR2#2: 5’- GCAGAGAACUUGAAAGCAUTT-3’, siTGFBR2#3: 5’- CCAUAUGCGGUGUGAAAUATT, siSMAD2: 5’-CAGGCCTTTACAGCTTCTCTT-3’, siSMAD3: 5’- GGCCATCACCACGCAGAACTT-3’, siSMAD4: 5’- GTACTTCATACCATGCCGATT-3’, siEZH2#1: 5’-GCUGACCAUUGGGACAGUATT-3′, siEZH2#2: 5’-GUGUAUGAGUUUAGAGUCATT-3′.

**Sequence of primers used for transcript analyses**

RT-MCM7-F: 5′-TCGAGGCATGAAAATCCGGG-3′

RT-MCM7-R: 5′-CGCCAGTCGATCAATGTATGACA-3′

RT-YAP-F:5′-CACAGCATGTTCGAGCTCAT-3′

RT-YAP-R: 5′-GATGCTGAGCTGTGGGTGTA-3′

RT-TAZ-F: 5′-CCATCACTAATAATAGCTCAGATC-3′

RT-TAZ-R: 5′-GTGATTACAGCCAGGTTAGAAAG-3′

RT-TEAD1-F: 5′-CCACCAAAGTTTGCTCCTTTGGGA-3′

RT-TEAD1-R: 5′-ACTTCAAACACACAGGCCATGCAG-3′

RT-GAPDH-F 5′-GAGTCAACGGATTTGGTCGT-3′

RT-GAPDH-R: 5′-GACAAGCTTCCCGTTCTCAG-3′

RT-EZH2-F: 5’-TGGTTAACGGTGATCACAGGA-3’

RT-EZH2-R: 5’-GGAGGTAGCAGATGTCAAGG-3’

RT-TGFBR2-F: 3’- GTAGCTCTGATGAGTGCAATGAC-5’

RT-TGFBR2-R: 5’- CAGATATGGCAACTCCCAGTG-3

RT-PUMA-F: 5’- GCAGGCACCTAATTGGGCT -3’

RT-PUMA-R: 5’- ATCATGGGACTCCTGCCCTTA-3’

RT-p15-F: 5’- GGACTAGTGGAGAAGGTGCG-3’

RT-p15-R: 5’-CATCATGACCTGGATCGCGC-3’

RT-p16-F: 5’- CCCCTTGCCTGGAAAGATAC-3’

RT-p16-R: 5’-AGCCCCTCCTCTTTCTTCCT-3’

RT-p21-F: 5′-GGGACAGCAGAGGAAGAC-3′

RT-p21-R: 5′-GCGTTTGGAGTGGTAGAAATC-3

RT-SMAD2-F: 5’- ACCGAAATGCCACGGTAGAA-3′

RT-SMAD2-R: 5- TGGGGCTCTGCACAAAGAT-3′

RT-SMAD3-F: 5′- GCCTGTGCTGGAACATCATC-3′

RT-SMAD3-R: 5′- TTGCCCTCATGTGTGCTCTT-3′

RT-SMAD4-F: 5′- CATCCTGCTCCTGAGTATTGG-3′

RT-SMAD4-R: 5′- GGGTCCACGTATCCATCAAC-3′

RT-EED-F: 5′-GTGACGAGAACAGCAATCCAG-3′

RT-EED-R:5′-TATCAGGGCGTTCAGTGTTTG-3′

RT-SUZ12-F: 5′-AGGCTGACCACGAGCTTTTC-3′

RT-SUZ12-R: 5’-GGTGCTATGAGATTCCGAGTTC-3’

RT-SMAD7-F: 5’-CTGCAGACTGTCCAGATGCTGTG-3’

RT-SMAD7-R: 5’-GGCTCCAGAAGAAGTTGGGAATCTGA-3’

RT-MYC-F: 5'-CTCCTGGCAAAAGGTCAGAG-3'

RT-MYC-R: 5'-TCGGTTGTTGCTGATCTGTC-3'

RT-HOXA5-F: 5′-TCTCGTTGCCCTAATTCATCTTT-3′

RT-HOXA5-R: 5′- CATTCAGGACAAAGAGATGAACAGAA-3′

RT-PTEN-F: 5′-AGTTCCCTCAGCCGTTACCT-3′

RT-PTEN-R: 5′- AGGTTTCCTCTGGTCCTGGT-3′

**Sequence of primers used for ChIp**

ChIP-CTGF-F: 5′-CTTTGGAGAGTTTCAAGAGCC-3′

ChIP-CTGF-R:5′-TCTGTCCACTGACATACATCC-3′;

EZH2Neg ctrl F: ChIP-EZH2-INTNEG-F: 5′- TCCTCTGGCATTTCTAGGGAATGTGGT-3′

EZH2Neg ctrl R: ChIP-EZH2-INT-NEG-R: 5'-

CAGACGTACTAAAATACAATATAGGAAAAA CCAAACT-3'.

ChIP-EZH2P1-F: 5'- GATGTCTCCCGGTCCCC -3'

ChIP-EZH2P1-R: 5'- TCTTTCGCTGAACACACGGC-3

ChIP-EZH2-P2-F: 5'-AGGCGT TCACCAAGTTTTCCAAAAGATCGT-3'

ChIP-EZH2-P2-R: 5' -CCCACTTCATTGTGT ACA TCC CCT TC-3'.

ChIp-TGFBR2-P1-F: 5'-CAGCTGAAAGTCGGCCAAAG-3'

ChIp-TGFBR2-P1-R: 5'-AGCCCCTAGCTCTCTCGTAG-3'

CHIP-TGFBR2ENHUP-F: 5'-CCT GAA GGT AAA AGT GGC ACA GAG TGT-3'

CHIP-TGFBR2ENHUP-R: 5'-CAA TTC ACA ACA GCT CTC TAA ATG CAG AAA CT-3'

CHIP-SMAD7UP-F: 5'-AAG AAG AAA CCA TTG AGG AGA CCA AAA AAG TAC-3'

CHIP-SMAD7UP-R: 5'-CTT TTC ATA AGC CAT CAC TTC CAT TTG TAT TGT TGC-3'

CHIP-CDKN1A-P-F: 5'-TGG GCT GCC TGT TTT CAG GTG AGG A-3'

CHIP-CDKN1A-P-R: 5'-TGC TGG CAG ATC ACA TAC CCT GTT CAG-3'

CHIP-PTENP2-F: 5'-AGCAAGCCCCAGGCAGCTACACT-3'

CHIP-PTENP2-R: 5'-GGTAGGAGGGGGCAGAGCGGTA-3'

ChIP-HOXA5-P1-F: 5'-TGG TAG TCC GGG CCA TTT GGA TAG C-3'

ChIP-HOXA5-P1-R: 5'-ATA GAC GCA CAA ACG ACC GCG AGC-3'

ChIP-CDKN2B-P2-F: 5'-GGC CCC AGC TAC CTG GAT CG-3'

ChIP-CDKN2B-P2-R: 5'-TCT GGG GGC TGC GGA ATG CG-3'

CHIP-P15-F: 5'-TTTGGCCTCCTCCCCAAATG-3'

CHIP-P15-R: 5'-ACGGCAGATGGGAATTCGTT-3'.
